# Hallucinating structure-conditioned antibody libraries for target-specific binders

**DOI:** 10.1101/2022.06.06.494991

**Authors:** Sai Pooja Mahajan, Jeffrey A. Ruffolo, Rahel Frick, Jeffrey J. Gray

## Abstract

Antibodies are widely developed and used as therapeutics to treat cancer, infectious disease, and inflammation. During development, initial leads routinely undergo additional engineering to increase their target affinity. Experimental methods for affinity maturation are expensive, laborious, and time-consuming and rarely allow the efficient exploration of the relevant design space. Deep learning (DL) models are transforming the field of protein engineering and design. While several DL-based protein design methods have shown promise, the antibody design problem is distinct, and specialized models for antibody design are desirable. Inspired by hallucination frameworks that leverage accurate structure prediction DL models, we propose the F_v_Hallucinator for designing antibody sequences, especially the CDR loops, conditioned on an antibody structure. Such a strategy generates targeted CDR libraries that retain the conformation of the binder and thereby the mode of binding to the epitope on the antigen. On a benchmark set of 60 antibodies, F_v_Hallucinator generates sequences resembling natural CDRs and recapitulates perplexity of canonical CDR clusters. Furthermore, the F_v_Hallucinator designs amino acid substitutions at the V_H_-V_L_ interface that are enriched in human antibody repertoires and therapeutic antibodies. We propose a pipeline that screens F_v_Hallucinator designs to obtain a library enriched in binders for an antigen of interest. We apply this pipeline to the CDR H3 of the Trastuzumab-HER2 complex to generate *in silico* designs improving upon the binding affinity and interfacial properties of the original antibody. Thus, the F_v_Hallucinator pipeline enables generation of inexpensive, diverse, and targeted antibody libraries enriched in binders for antibody affinity maturation.

## Introduction

Antibodies recognize and bind an extremely large repertoire of antigens *via* six hypervariable loop regions (H1, H2, H3, L1, L2, L3) in their variable domain (F_v_) known as the complementarity determining regions (CDRs). The CDRs leverage a vast sequence space to target the immensely diverse range of antigens that challenge the immune system. CDR diversity results from V(D)J gene recombination prior to antigen exposure followed by somatic hypermutation after antigen exposure.[1] Hence, antibodies achieve the ability to target a diverse range of epitopes by both diversifying the CDR sequences prior to antigen exposure and by antigen-specific hypermutation post antigenic exposure. The process of evolution of an antibody to bind an antigen with higher affinity and specificity is known as affinity maturation.

In the laboratory, affinity maturation is achieved broadly in two steps. First, large libraries of CDR regions are diversified *via* methods such as random mutagenesis, targeted mutagenesis, and chain shuffling. Second, the libraries are screened for expression and binding through display technologies such as yeast or phage display. These steps are repeated until enough “hits” are found with the desired affinity. Such approaches to affinity maturation can be expensive and time-consuming, and rarely allow the efficient exploration of the full design space.[2] Computational methods offer faster and inexpensive alternatives to experimental affinity maturation. Conventional computational approaches for antibody design or affinity maturation include rational or structure-guided design strategies[3,4], general protein design methods such as FastDesign[5], and antibody-specific design methods such as AbDesign[6] and RosettaAntibodyDesign[7] (RAbD). RAbD is notable because it allows the design of CDR sequences and conformations in the context of the antigen. However, RAbD requires 10-20 hours for a single design. Further, RAbD only samples CDR sequences from PyIgClassify[8] clusters that have arbitrary classification cutoffs and are context (surrounding residues) agnostic. Further, Rosetta (like other methods) has challenges in accurate modeling of CDR H3.[9]

Deep learning (DL) models are transforming the field of protein structure-prediction, engineering, and design.[10–12] Over the last few years, DL models have emerged as the leaders in predicting protein structures with high accuracy, and they are increasingly being applied to protein design.[11,13,14] For the purpose of protein design, DL models fall in three broad categories, 1) Sequence generation with language models[15,16] 2) Structure-conditioned sequence generation[17,18], and 3) Sequence agnostic structure or backbone generation[19,20]. Since the antibody design task is primarily focused on CDRs that are regions of high variability and flexibility, it may require specialized DL models.[21]. An example of an antibody-specific DL model is IgLM, a language model that generates variable-length CDR sequence libraries conditioned on chain type and/or species-of-origin.[22] IgLM designed synthetic libraries are akin to naïve libraries that can be further screened to obtain a lead antibody sequence. Another antibody-specific DL model treats the problem of antibody CDR generation as an iterative sequence-structure prediction problem.[23] It also proposes a sequence-based affinity maturation protocol that conditions design on known sequences of binders against a target antigen. This approach is promising when a sufficiently large library of sequences that bind an antigenic epitope is available.

Here, we propose a fast and versatile DL framework for antibody design and engineering that is aimed at shortening the cycle of antibody library generation and affinity maturation. Given that structural information is becomingly increasingly abundant and accessible [11,24], our framework (like RAbD[7]) leverages structural information of both the antibody and the antigen.

Our approach is inspired by “hallucination frameworks,” a special class of DL based design methods, that leverage highly accurate pretrained sequence to structure models for protein design.[12,25,26] Hallucination frameworks have been used to design *de novo* sequence-structure pairs,[12] sequences that maximize the likelihood of a target structure,[25] and protein scaffolds that can host functional motifs[26].

We adapted the trDesign[27] approach for the specific task of generating antibody libraries conditioned on a target antibody structure. Our framework (Figure 1A), F_v_Hallucinator, differs from the previous hallucination frameworks in three important aspects. First, it is developed for the variable domains of conventional antibodies (F_v_ region). To estimate the likelihood of a structure given a sequence, we use an antibody specific sequence-to-structure model, DeepAb[28]. DeepAb improves the prediction of CDR H3 loop structure over conventional approaches such as RosettaAntibody.[28] Second, the framework can be applied to hallucinate sequences that optimize the heavy and light chain interface[29], a task that is not suitable for hallucination frameworks with models trained on single sequences (trRosetta[30], RoseTTAFold[31]). And lastly, though the framework is applicable to the design of any subset of residues in the F_v_ region, its main purpose is to generate libraries of CDRs, the hypervariable loop regions that bind the antigen. In this regard, the F_v_Hallucinator is distinct from and complements existing hallucination approaches that either design full proteins that fold into a target structure or scaffolds that host fixed functional motifs.

**Figure 1.**
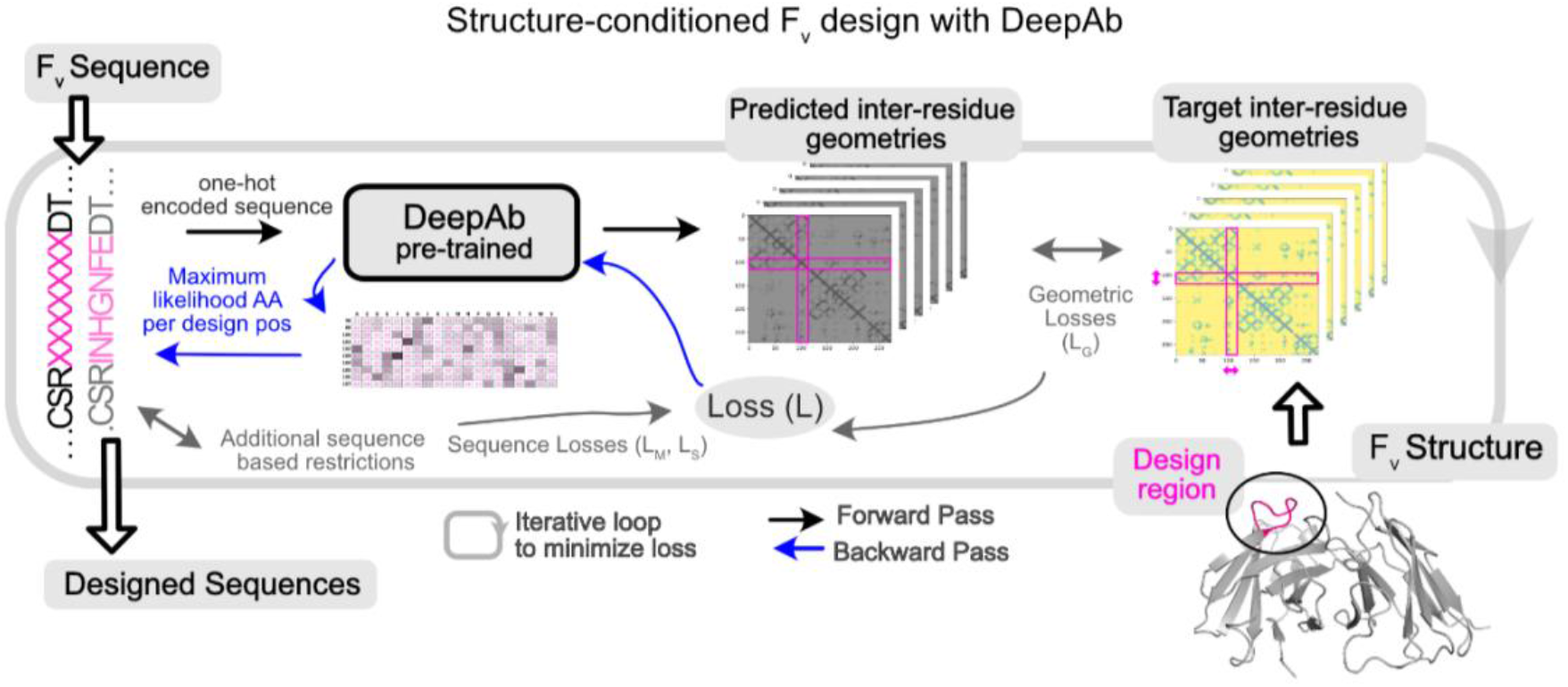
Hallucination framework (F_v_Hallucinator) for generating antibody F_v_ libraries conditioned on structure. F_v_Hallucinator is adapted from trDesign[27] for antibody design. An ensemble of pretrained DeepAb[28] models is used to predict the structure of a designed sequence. The error/loss between predicted structure and target structure is minimized iteratively to arrive at a sequence that folds into the target structure as predicted by DeepAb.

The F_v_Hallucinator aims to achieve targeted, fast, high-throughput and combinatorial sampling of the CDR sequence space while maintaining the conformation of the original binder. To further select antigen-specific designs, we propose a pipeline to virtually screen the hallucinated sequences against a target antigen. This pipeline screens for sequences that retain binding to the antigen in the same mode as the original binder from a large and diverse structure-conditioned hallucinated library. We show that such targeted exploration of the sequence space enables *in-silico* improvements in binding energies and other desirable antibody-antigen interface metrics such as hydrogen bonding, shape complementarity and buried surface area at the interface.

## Results and Discussion

### Structure-conditioned subsequence generation

#### A framework for F_v_ hallucination

We aim to design sequences that fold into a desired F_v_ structure by leveraging a pretrained sequence-to-structure prediction DL model. We adapt the trDesign[27] approach where the problem of predicting sequence given structure has been reframed as the problem of *maximizing the conditional probability of a sequence given structure*. In the case of the F_v_, we are primarily interested in designing a subset of the residues (CDRs, V_H_-V_L_ interface), so we split the sequence *S* into fixed and designable positions, *S_F_* and *S_D_*. We then seek the design subsequence *S_D_* that maximizes the conditional probability of the sequence *S* given a target structure *T* and the fixed sequence *S_F_*:

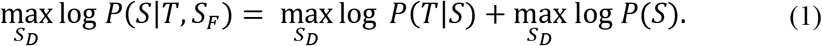

The log *P*(*T*|*S*) is maximized by minimizing the categorical cross entropy loss of the DeepAb model (or geometric loss; *L_G_*), for sequence *S* and structure *T*, with respect to the design subsequence *S_D_*. The log *P*(*S*) term is constant for hallucination guided only by geometric losses or, to sample sequences biased toward a particular sequence or a motif, maximized by minimizing a sequence-based loss. Alternatively, we may alter *P*(*S*) by initializing the design subsequence with an increased likelihood for a sequence (*e.g.*, “Wildtype Seeding” for seeding initial sequence with the wildtype sequence) or by initializing from a subset of the amino acid alphabet (e.g., omitting sampling of a particular amino acid at design positions at the time of sequence initialization). For the full formulation, see Methods section.

**Figure 1** shows the workflow architecture. First, we randomly initialize subsequence *S_D_* from the amino acid alphabet. Second, we input the full sequence to the DeepAb model to predict inter-residues distances and orientations (geometries), a proxy for structure. Third, we calculate the geometric loss (see methods) between the predicted structure representation and the target structure representation and, in some cases, additional non-geometric losses for restricting designs to relevant sequence spaces (detailed later). Fourth, to minimize the geometric loss, we calculate the gradient of the loss with respect to the design subsequence (*S_D_*) with the Stochastic Gradient Descent (SGD) optimizer and normalize the gradient[25], resulting in an updated subsequence matrix (number of design positions x 20) that represents the probability of each amino acid for each design position. Finally, we reduce this subsequence matrix to a sequence (one amino acid per design position) by choosing the amino acid with the maximum probability at each design position. This designed sequence is input back to the model by returning to step two. We repeat steps two to five until the loss is reduced to a small value to yield a designed sequence that folds into the target structure. Forty to sixty iterations are required to reach convergence for a single CDR design (SI Figure 1). A single sequence design requires under 3 mins on an NVIDIA A100 GPU, and several designs can be generated in parallel. All design runs (50-1200 designs per task) reported in this work were accomplished in a few hours with little or no parallelization (1-5 GPUs). The framework (**Figure 1**) is fully automated, highly versatile, and customizable for different design objectives mostly implemented through different priors for the design sequence (*P*(*S*) in Equation 1).

### Structure-conditioned sequence design of CDRs and the V_H_-V_L_ interface

#### Hallucination recovers native-like sequences on benchmark set of 60 antibodies

To test whether the F_v_Hallucinator can recover native-like CDR sequences corresponding to the structures it was conditioned on, we measured the amino acid sequence recovery (AAR) for each CDR loop on a benchmark set of 60 antibodies first introduced in the RAbD study.[7] Fifty sequences were designed each for CDRs H1, H1, L1, L2 and L3, and 100 sequences for CDR H3 (see Methods). We calculated AAR as the percentage of native residues recovered per designed CDR averaged over all 50 designs.

We expect the AARs to be limited since CDRs are surface-exposed and evolve in the context of the antigen. Rotamer packing methods such as Rosetta report an AAR of less than 27% for surface residues.[32] Indeed, the probability of recovering the full wildtype sequence even when sampling directly from position specific scoring matrix of PyIgClassify cluster that corresponds to its target structure is vanishingly small (SI Figure 2). However, all CDR loops except H3 fold into a small number of “canonical structures” characterized by structural motifs conferred in part by a few key residues[33]. Therefore, we expect hallucinated sequences to recover the residues that are crucial for target structure realization. Moreover, if the algorithm is seeded to search in the vicinity of the native sequence, we expect to achieve higher AARs as the native residues will be retained with higher probability since the native sequence folds into the target structure. To this end, we also performed hallucination with “wildtype seeding” where the starting sequence of the design region is sampled using wildtype residue types with higher probability than random (see Methods).

**Figure 2A** shows the sequence recovery on all six CDRs for the benchmark set conditioned on the native CDR loop conformation with (dark blue) and without (light blue) wildtype seeding. Without wildtype seeding, AAR is lower because the algorithm recovers only the more conserved residues (see SI Figure 3 for AAR for all, top 50% most conserved and top 30% most conserved residues). With wildtype seeding, the algorithm recovers over 50% of the wildtype residues. The AARs for all 6 CDRs with wildtype seeding is competitive with 50-70% AAR reported for RAbD (initialized with the wildtype sequence and amino acid sampling restricted to PyIgClassify cluster motif of the wildtype CDR when sampled without the antigen)[7].

**Figure 2.**
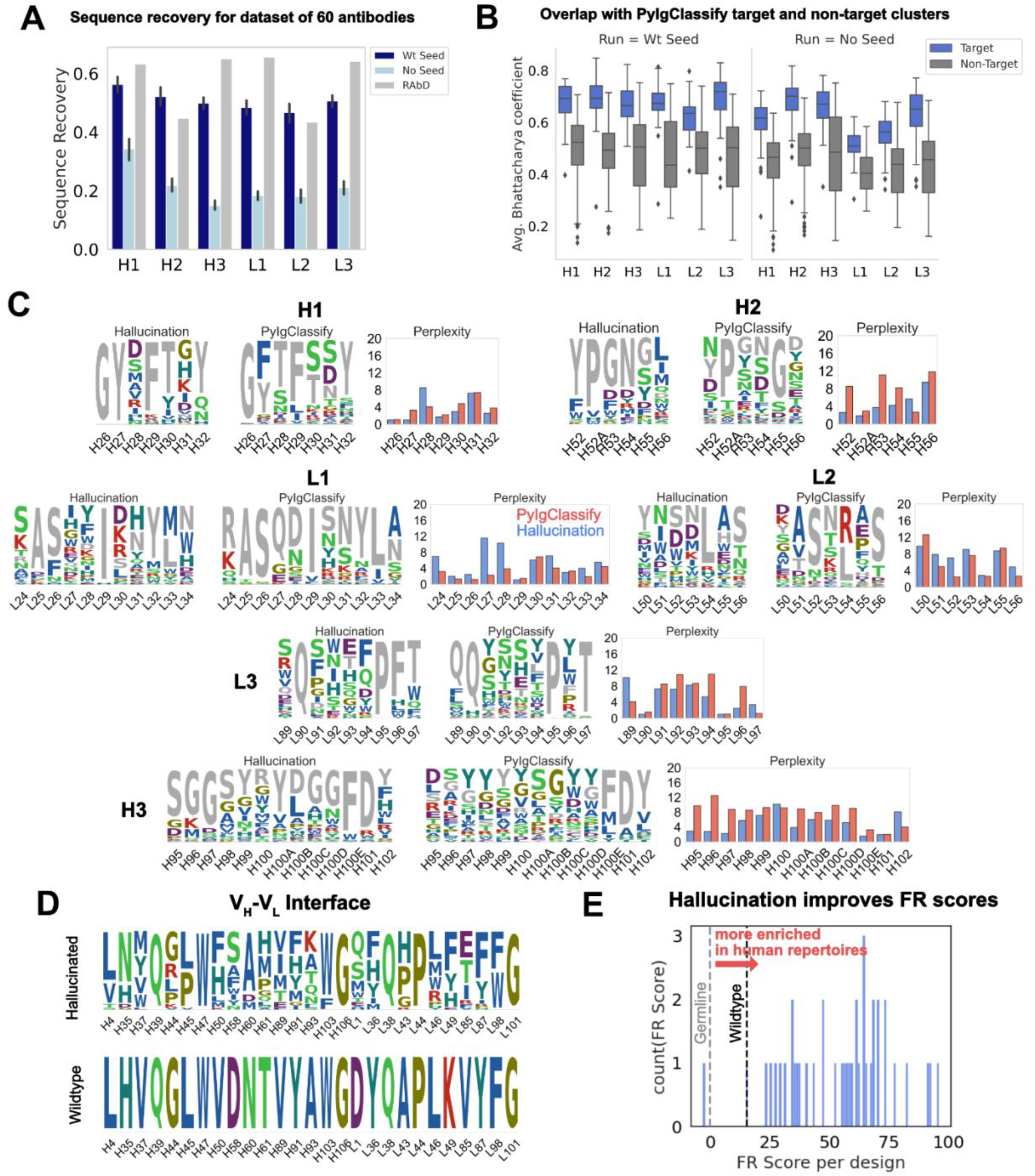
Hallucinated, structure-conditioned sequence libraries for CDR loops and the V_H_-V_L_ interface. (A) Sequence recovery on the 60 antibody RAbD benchmark set [7] from hallucination with and without wildtype seeding. RAbD values as reported in [7] for sequence recovery without antigen. (RAbD sequence recovery is reported separately for contact and non-contact residues. To obtain a single value per-CDR, we obtained the weighted average of contact and non-contact sequence recovery, weighted by the number of contact and non-contact residues for each CDR over the full dataset. Also see SI Figure 3 for AAR for all, Top 50% most conserved and Top 30% most conserved residues.) (B) Average Bhattacharya coefficient between hallucination sequence profiles and PyIgClassify[8] sequence profiles of target and non-target clusters (with and without wildtype seeding). Averages are over all positions on the CDR. (C) Comparison of distribution perplexity of hallucinated sequence profiles for CDRs with PyIgClassify profile of the corresponding CDR cluster for anti-neuraminidase influenza virus antibody[34] (PDB ID: 1A14). Wildtype CDR sequence is colored in grey. Profiles were generated with wildtype seeding (SI Figure 12 shows the same for hallucinated profiles from runs without wildtype seeding.) (D) Hallucinated sequence profile for the V_H_-V_L_ interface for humanized antibody hu225 [37] (*no seeding*). (E) Distribution of FR scores for V_H_-V_L_ interface designs shown in (D). A higher FR score signifies amino acid residues with higher enrichment in human repertoires.

To benchmark hallucination on a blind dataset, we also obtained sequence recovery on 20 antibodies (SI Table 1) selected from the DeepAb test set[28]. The average sequence recoveries are comparable to those obtained for the RAbD dataset (SI Figure 4).

#### Designed sequences’ distribution overlaps that of the target canonical cluster

To test whether the hallucinated sequence profiles exhibit distributions similar to those of the known sequences that fold into the target conformation, we obtained sequence profiles from the PyIgClassify[8] database of CDR structure clusters. To measure the overlap between the hallucinated sequence profiles and the PyIgClassify sequence profiles, we calculated the average Bhattacharyya coefficient (see Methods) over all positions on the CDR loop between the hallucinated sequence profiles and the PyIgClassify sequence profile of the target cluster and non-target clusters (**Figure 2B**; SI Figure 5). For all CDRs, the average Bhattacharya coefficient is about 0.2 higher for the sequence profiles belonging to the target structure cluster than the non-target clusters i.e., hallucinated sequence profiles have higher overlap with the known sequences that fold into the target cluster. This trend is observed for each target antibody (see SI Figure 6-10 for BC for each target for all six loops). Although longer CDR H3 loops cannot always be categorized into well-resolved clusters due to higher sequence and structural diversity, in cases where there are multiple PyIgClassify clusters available for the target CDR H3, the hallucinated profiles have a higher overlap with the target cluster than the non-target clusters (SI Figures 6-10).

#### CDR designs retain conserved sequence motifs and exhibit diversity

As an example of the CDR sequence profiles generated from hallucination (with wildtype seeding), **Figure 2C** shows the sequences sampled for all six loops of the anti-neuraminidase influenza virus antibody[34] (PDB id: 1A14; RAbD dataset) as sequence logos. (SI Figure 11 for effect of wildtype seeding on the hallucinated sequence profile and SI Figure 12 for sequence logos without seeding). We juxtapose each sequence logo with the sequence logo for the PyIgClassify cluster corresponding to the target CDR loop structure. To compare whether hallucination correctly captures the conservation and diversity observed in native sequences given a target conformation, we compared the perplexity at each position on the CDR of the hallucinated profiles to those of the PyIgClassify target cluster (**Figure 2C**). Several low (H26, H29, H32, H52A, H101, L25, L26, L29, L54, L90, L95) and high (H31, H100, L30, L34, L53, L55, L91, L93) perplexity positions in canonical clusters are recapitulated in hallucinated profiles (also see SI Figure 12). However, at certain positions, hallucination perplexities differ from PyIgClassify. Hallucination suggests that positions H28, L27, and L28 can be varied more than the PyIgClassify profiles suggest, and conversely hallucination does not capture the diversity of observed profiles at positions H52, H53, H95, H96, and L96. In the case of the H3 residues (H95, H97, H97), hallucination suggests that the wildtype residues are more important to the H3 conformation than the PyIgClassify profile suggests. Also, some discrepancies between hallucinated and PyIgClassify profiles are expected as both DeepAb and PyIgClassify profiles are only estimates of a large and incompletely mapped sequence-structure space.

As a control, we hallucinated designs with wildtype seeding but with scrambled geometric losses. These experiments showed that incorrect geometric losses generate random sequence sampling (SI Fig. 11). In a second control we removed the wildtype seeding. These experiments showed increased perplexities (that is, greater variation in designs) for most but not all positions (SI Fig. 12). Several key conformation-determining positions retained critical motif residues (e.g., much of H1), and the H3 perplexities were similar to those of PyIgClassify.

#### Hallucinated V_H_-V_L_ interface designs accumulate mutations enriched in human repertoires and therapeutic antibodies

Improvements in both stability and affinity can be achieved by optimizing the V_H_-V_L_ interface[29]. Hence, we applied the F_v_Hallucinator framework to design non-CDR V_H_-V_L_ interface residues. Another important consideration in evaluating non-CDR designs is the likeness to human antibodies. Since humanization of mouse antibodies remains the primary route to therapeutic antibody development, a design method that improves the humanness of antibodies while retaining original contacts and structure is desired.

To evaluate the human-likeness of the hallucinated designs, we turned to the work of Petersen *et al.* [35], which investigated the amino acid preference for framework mutations in human repertoires and FDA approved therapeutic antibodies. Petersen *et al.*[35] calculated the frequency of amino acids in human repertoires at framework positions for 25 VH genes that represent the precursors of several FDA approved antibodies and many of the most common VH genes. They converted these frequencies into “FR scores”, a measure of amino acid enrichment over the germline residue at each framework position. They found that mutations with higher FR scores result in lower immunogenicity and are highly enriched in FDA approved antibodies. [35] Hence, we calculated the FR scores to evaluate the humanness of hallucination-designed mutations at the V_H_-V_L_ interface.

To evaluate the potential of hallucination in improving the humanness of the framework residues at the V_H_-V_L_ interface, we collated a set of nine humanized antibodies and designed their V_H_-V_L_ interfaces. A significant percentage of designs exhibited FR scores greater than the wildtype (Table 1, SI Figure 13). **Figure 2D** shows the sequence profiles of the V_H_-V_L_ interface designs for the antibody hu225, a humanized version of the mouse-derived therapeutic antibody Cetuximab[36]. **Figure 2E** shows the distribution of net FR scores per design (summed over 12 design positions) for 40 designs and the net FR score for the wildtype for comparison. These data show that hallucinated sequences have mutations that are preferentially accumulated in human repertoires during antibody maturation, suggesting that hallucinated antibodies are more human-like and less immunogenic than the starting antibody [35].

**Table 1.**
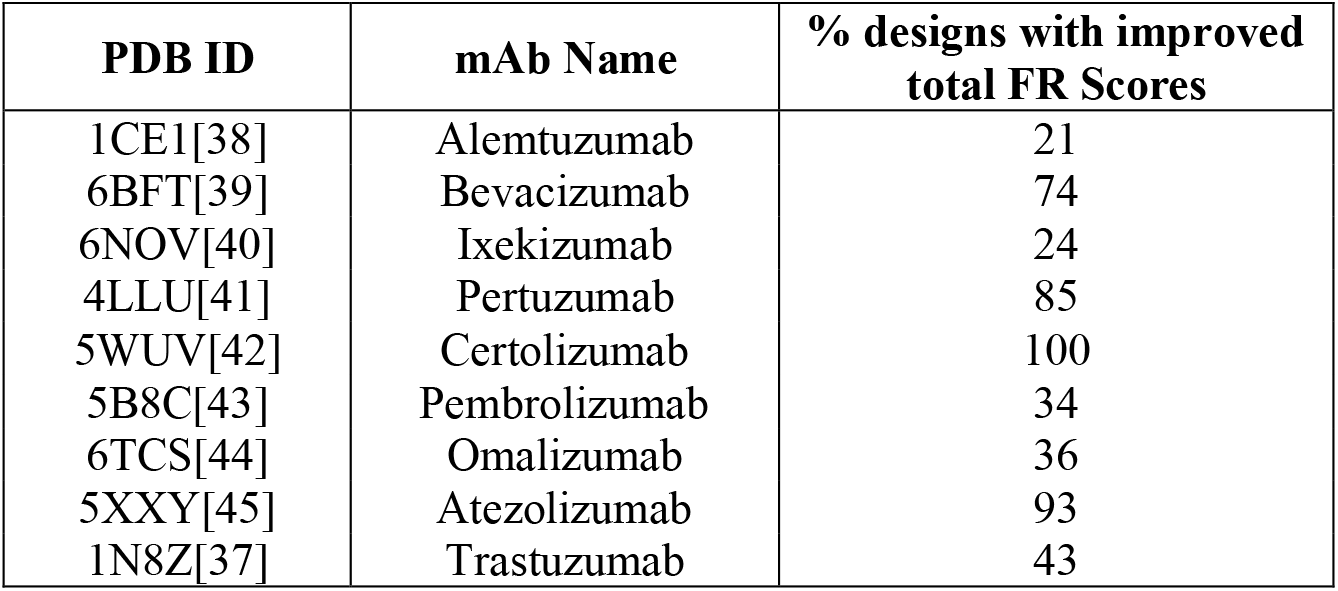
Percentage of V_H_-V_L_ hallucinated designs with improved FR scores (over wildtype) for a selected set of 9 humanized therapeutic antibodies. A total of 60 designs were obtained for each case with no seeding. See SI Figure 13 for distribution of FR scores per target.

| PDB ID | mAb Name | % designs with improved total FR Scores |
| --- | --- | --- |
| 1CE1[38] | Alemtuzumab | 21 |
| 6BFT[39] | Bevacizumab | 74 |
| 6NOV[40] | Ixekizumab | 24 |
| 4LLU[41] | Pertuzumab | 85 |
| 5WUV[42] | Certolizumab | 100 |
| 5B8C[43] | Pembrolizumab | 34 |
| 6TCS[44] | Omalizumab | 36 |
| 5XXY[45] | Atezolizumab | 93 |
| 1N8Z[37] | Trastuzumab | 43 |

### Sequence losses to restrict hallucination to relevant sequence spaces

Many CDR sequences can fold into the same conformation. So, when sampling with the geometric loss, the solvent exposed residues of a CDR will sample a large and unrestricted sequence space. Unrestricted hallucination, guided solely by DeepAb’s geometric loss, is apt when the goal is to sample a large and diverse sequence space only restrained by loop conformation. However, sometimes we seek designs close to a known sequence, for example to retain core antigen-binding residues (**Figure 3A**). To address such design objectives, we developed two restricted modes of hallucination. In sequence-restricted hallucination, we apply a sequence loss to sample amino acid residues close to a given sequence. In motif-restricted hallucination, we apply a motif loss to sample amino acid residues in specified proportions at specified positions (e.g., 50% Y and 50% S at position 100A on the heavy chain). These sequence-based losses are added to the geometric loss during backpropagation to update the sequence at the design positions (**Figure 3B**, see Methods).

**Figure 3.**
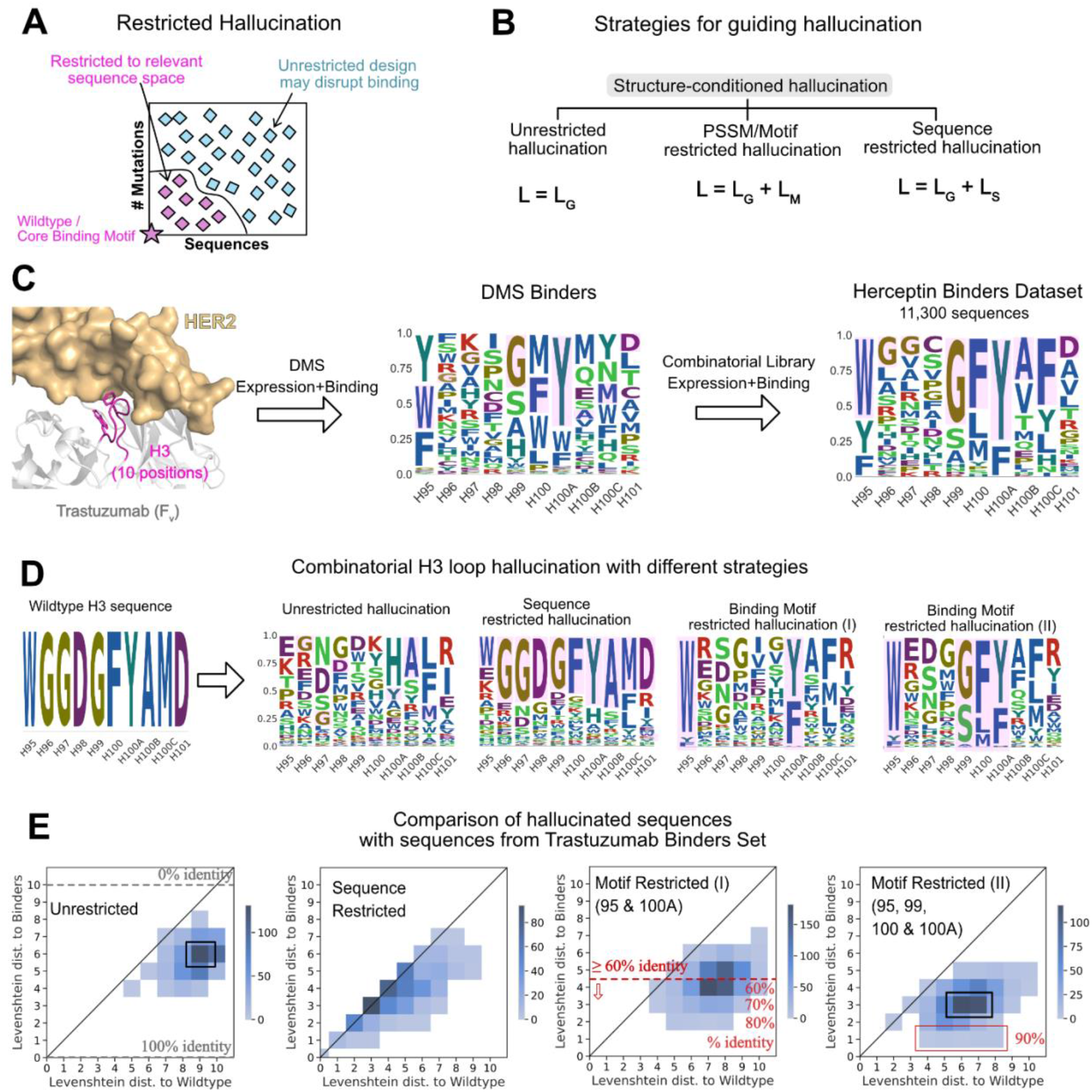
Strategies for guiding hallucination to relevant sequence spaces. (A) Restricted hallucination allows restricting hallucination to relevant sequence spaces for specific design objectives such as sampling close to the wildtype sequence or retaining motifs for binding. (B) Guiding hallucination towards different sequence spaces via geometric losses (*L_G_*, unrestricted hallucination), geometric and motif loss (*L_M_*, motif restricted hallucination), and geometric and sequence loss (*L_S_*, sequence restricted hallucination). (C) Trastuzumab binders dataset (TBS) from Mason *et al.* [46] screening of single-site and combinatorial libraries for expression and binding. (D) Hallucinated sequences shown as sequence logos for 3 modes of hallucination for the CDR H3 loop of Trastuzumab. (E) Comparison of LDs of hallucinated sequences with different strategies with experimentally identified binders from the TBS dataset.

#### Binding-motif restricted designs are consistent with experimentally screened binder sequences

To compare hallucinated designs to experimentally generated CDR libraries, we chose a dataset of 11,300 unique CDR H3 sequences (Trastuzumab Binder Set; TBS). To obtain this dataset, Mason *et al.* [46] first constructed a single-site deep mutational scanning (DMS) library for the CDR H3 of Trastuzumab (trade name Herceptin) and screened them in multiple rounds of expression and binding to HER2. This resulting library was highly restricted at 4 positions, namely, H95, H99, H100 and H100A (Chothia numbered) to F/W/Y, G/S/A, F/L/M and Y/F, respectively, while the remaining positions varied. Mason *et al.* then combined this profile to generate a combinatorial library and screened it in multiple rounds for expression and binding to HER2. Next generation sequencing of the binding population from the final round yielded 11,300 unique CDR H3 binders (**Figure 3C**) that were highly diverse at 6 out of 10 positions, suggesting that the CDR H3 may retain binding to the antigen with little to no overlap with the wildtype sequence.[46]

To compare hallucinated designs to the TBS, we generated designs in three separate modes – unrestricted hallucination, sequence-restricted hallucination (with wildtype as the target sequence) and with two different motif-restricted hallucinations. **Figure 3D** shows the sequence profiles of the designs generated from each mode. In **Figure 3E**, for each run, we show the joint distribution of the minimum Levenshtein distance (LD) of the designed sequences to the TBS sequences and the LD of the same to the wildtype sequence.

With unrestricted hallucination, the largest fraction of designs (**Figure 3E**, black box) exhibit only 10% identity (LD 9) with the wildtype yet recover 40% sequence positions of one or more binders in the TBS. Additionally, a small fraction of sequences exhibit about 50% sequence identity (LD 5) with the wildtype sequence and 60% identity (LD 4) with one or more binders.

Sequence restricted hallucination samples a sequence space that largely retains the wildtype sequence. Most designs are equidistant from the wildtype sequence and one or more binders (**Figure 3E**).

Since the experimental library was generated with each position restricted to the relative fractions of amino acid residues in the DMS profile, motif-restricted hallucination is the most comparable mode of hallucination. For the motif-restricted hallucination, where we restricted only two (H95 and H100A to F/W/Y and Y/F respectively) positions that form the core-binding motif between CDR H3 and the antigen, about 50% of hallucinated sequences show over 60% identity with one or more binders (**Figure 3E,** red line). A small fraction of hallucinated sequences exhibits 80% sequence identity (**Figure 3E**) with a binder. As expected, when we restricted all four positions that were restricted in the DMS library (H95, H99, H100 and H100A to F/W/Y, G/S/A, F/L/M and Y/F, respectively), the overlap with TBS increases (Figure 3E, rightmost panel). However, the maximum sequence identity obtained with four restrictions is 90% (**Figure 3E**, red box), considerably higher than the 40% sequence identity conferred just from restricting the four positions. Thus, restricting hallucination to relevant sequence spaces yields virtual libraries that have significant overlap with experimental libraries and is a useful strategy to generate structure-conditioned libraries tailored towards a desired design objective.

### A pipeline for screening antigen-specific sequences from hallucinated libraries

To enrich the hallucinated library in antigen-specific binders and to select for desired properties such as hydrogen bonding and high shape complementarity at the interface, the antigen-antibody structure can help. We propose a pipeline (**Figure 4A**) that first hallucinates a large library of structure-conditioned antibody sequences with or without additional restrictions. Next, we forward fold the designed sequences with DeepAb to validate that the sequences fold into the target structure, resulting in a structure-conditioned, antigen-agnostic library. Then, we virtually screen the library for antigen binding by refining a model antibody-antigen complex using Rosetta (based on the wildtype crystal structure) and measuring the free energy of binding to the antigen with Rosetta’s InterfaceAnalyzer [47,48]. Finally, we obtain the screened library by selecting the subset of designs (screened library) that satisfies both folding and binding thresholds. We characterize the screened library for various interface metrics to identify top designs for experimental characterization.

**Figure 4.**
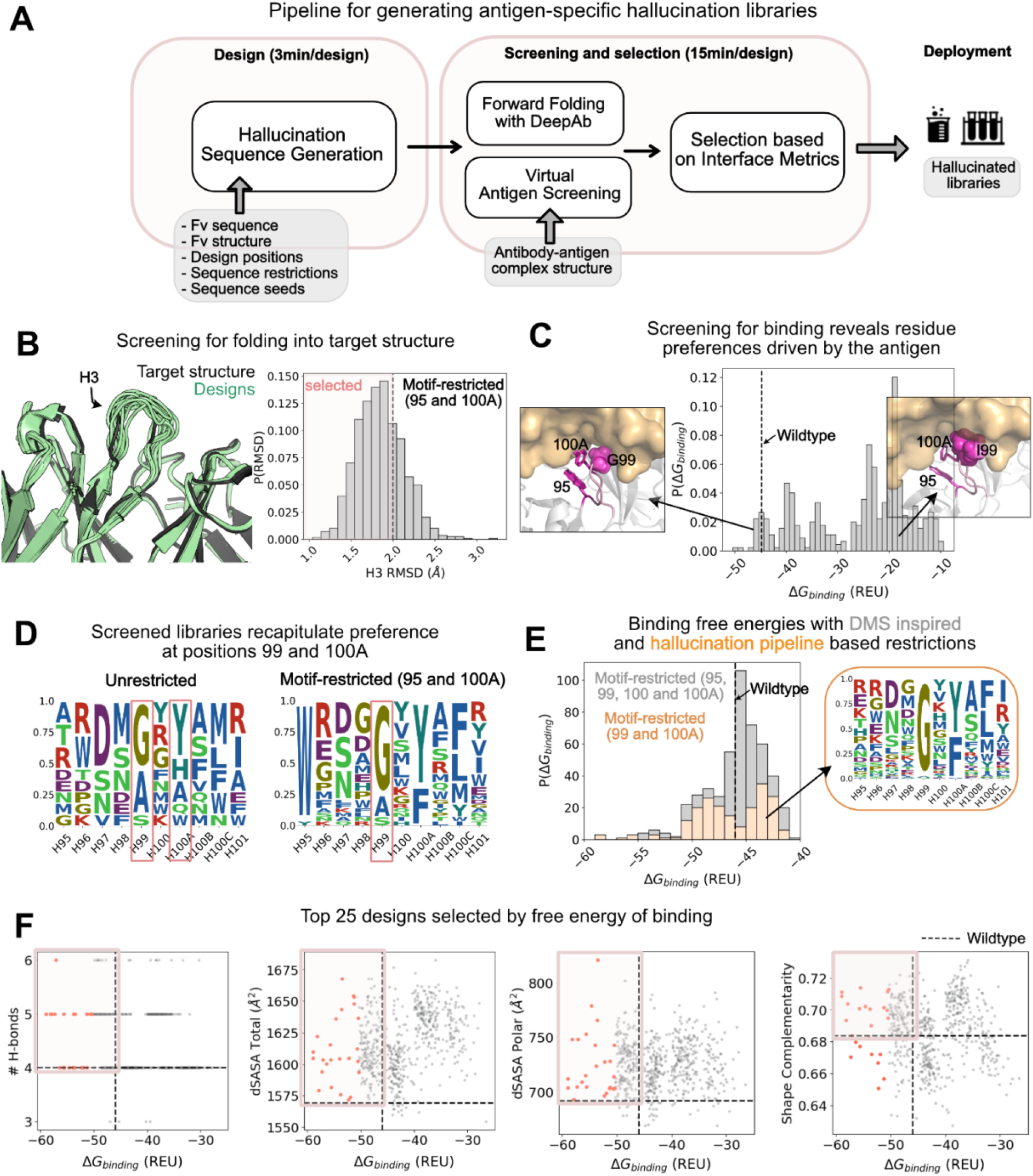
Application of hallucination pipeline to generate large number of unique designs with improved binding as characterized by various interface metrics. (A) A pipeline for screening the hallucinated library for folding into the target structure and binding the target epitope. The screened library is characterized *in silico* for interfacial metrics to select designs for experimental testing. (B) Comparison of structure of forward folded hallucinated sequences with wildtype structure: (Left) Randomly selected forward folded structures of hallucinated designs (green) with DeepAb superposed with wildtype structure (black). (Right) The distribution of the H3 *RMSD* of the designs with respect to the wildtype. Dashed line at *RMSD* = 2 Å marks the threshold for selecting designs that fold into the wildtype structure. *RMSDs* were calculated on all heavy atom backbone residues (N, CA, C) excluding (O) consistent with *RMSD* reported in DeepAb[28]. Distribution is shown for the motif-restricted hallucination I (95, 100A restricted) from Figures 3D and 3E. (C) Distribution of screened binding free energies against HER2 for the designs. The dashed line is the binding free energy of the wildtype Trastuzumab antibody. *Popouts: (Left)* Representative design with G at position 99 that exhibits binding energies better than or comparable to the wildtype. *(Right)* Representative design with an I at position 99 that exhibits binding energies significantly worse than the wildtype. (D) Sequence logos of *screened libraries* for the unrestricted hallucination and motif-restricted hallucination I (95 and 100A restricted). The unrestricted designs show strong preference for residues G and Y at positions 99 and 100A respectively. The motif-restricted (95 and 100A) mode additionally confirms the preference for Gly at position 99. Both restrictions (at positions 99 and 100A) are also observed in experiments. (E) Comparison of distributions of binding free energies for designs from two motif-restricted hallucinations: (gray) Motif-restricted hallucination with positions 95, 99, 100, 100A restricted to experimentally observed preferences, and (orange) motif-restricted hallucination with positions 99 and 100A restricted to hallucination-pipeline derived preferences from (D). *Popout:* Sequence logo for the screened library for designs from the latter hallucination. (F) Interface metrics for screened designs from motif-restricted hallucination (99 and 100A restricted). A significant fraction of designs exhibit metrics better than the wildtype (boxed quadrant); top 25 designs (selected by binding free energy) highlighted in orange.

We applied the full pipeline to generate a library of CDR H3s (positions 95 - 101) for the Trastuzumab antibody enriched in HER2 binders. We applied the pipeline to hallucinated libraries obtained from different hallucination modes described in the previous section and in **Figures 3D and 3E**.

#### Over 70% of the designs retain target conformation when forward folded with DeepAb

To screen the hallucinated designs for folding, we forward folded the designed sequences with DeepAb and measured the RMSD of the CDR H3 loop with respect to the Trastuzumab antibody structure used to condition hallucination. For the motif-restricted hallucination with positions 95 and 100A restricted, 433 of 600 hallucinated sequences retained conformations with CDR H3 RMSD ≤ 2.0 Å (**Figure 4B**). For almost all modes of hallucination explored in **Figures 3D and E**, over 70% of the hallucinated designs retained target conformation (CDR H3 RMSD ≤ 2.0 Å; SI Figure 14). We also measured the per-residue RMSD for the designed residues and found that the solvent exposed residues show the largest deviations, whereas the stem residues show little deviation from the target conformation (SI Figure 15). This is also apparent from the superposition of forward folded designs (pale green) with the wildtype antibody (black) as shown in **Figure 4B**. Hence, hallucination enables the exploration of a diverse sequence space distinct from the wildtype sequence, while also retaining a conformation close to the wildtype structure.

#### Virtual screening recapitulates restrictions at positions 99, 100A

To screen the hallucinated, structure-conditioned library for antigen binding, we measured the designs’ free energy of binding (*ΔG*_binding_) with Rosetta’s InterfaceAnalyzer [48]. In **Figure 4C**, we show the distribution of *ΔG*_binding_ for hallucinations with positions 95 and 100A motif-restricted (SI Figure 16 shows the distributions for sequences from other modes of hallucination). Only a small fraction of hallucinated designs exhibit a *ΔG*_binding_ comparable to or better than the wildtype.

To investigate why only a small fraction of hallucinated designs pass virtual screening, we analyzed the sequences of designs with favorable (relative to the wildtype) and unfavorable binding energies. Compared to the hallucinated sequence profiles (**Figure 3D**), the screened libraries are more restricted especially at positions 99 and 100A (**Figure 4D**). For the unrestricted mode of hallucination, the only mode where position 100A was not restricted, we also found that about 50% of the sequences screened for binding sampled a Tyr at 100A (**Figure 4D**), recapitulating experimentally observed preference for Tyr at this position. Furthermore, for designs from all modes of hallucination where position 99 wasn’t explicitly restricted (**Figure 4D** and SI Figure 17), sequences with favorable binding energies preferred Gly (and other smaller residues to some extent) at this position—a preference which also aligns with experimentally identified binders (**Figure 3C**).

This narrowing of preference at positions 99 and 100A is antigen-driven as the hallucinated library (antigen-agnostic) sampled a broad range of amino acid residues at these positions (**Figure 3D**) whereas the antigen-screened library reduced these positions to a small set of amino acid residues (**Figure 4D**).

To understand the structural basis of this preference, we visualized the structures of favorable designs (almost always with Gly) and unfavorable designs (without Gly) at position 99 (**Figure 4C**). The substitution of Gly with other residues (especially larger residues such as Ile) resulted in steric hindrance pushing the antigen away from the paratope leading to significant loss in binding energy. Here, the Rosetta binding energies proved to be useful in deciphering the structural context for the preference of Gly at position 99, showing that it can be advantageous to combine physics-based approaches such as Rosetta with DL frameworks.

#### Pipeline reveals a less restricted sequence space than experiments, amenable to further improvement in affinity

In **Figure 4D**, we show the sequence logos of the screened libraries for the unrestricted and motif restricted (positions 95 and 100A restricted) modes of hallucination. The screened sequences show amino acid restrictions primarily at positions 99 and 100A, suggesting that the CDR H3 may be able to retain binding to HER2 with fewer restrictions (two) than those inferred from single-site DMS experiments (four). One possible explanation is that while DMS explores a single-site substitution space, scanning one position at a time and retaining the wildtype sequence at all other positions, hallucination explores a combinatorial space designing all positions simultaneously constrained only by geometry (and or sequence/motif). Hence, DMS may limit the sequence space too close to the starting sequence and miss out on a potentially larger sequence space available to the CDR H3 that may be amenable to binding HER2.

To test this hypothesis computationally, we hallucinated designs with restrictions at positions 99 and 100A only. Next, we calculated the binding free energy for this less-restricted library (**Figure 4E**, orange bars). Remarkably, this scheme of hallucination yielded a library that was highly enriched in binders. Over 50% of the designs exhibited binding energies comparable or better than the wildtype (dashed line). Moreover, 27% designs exhibit lower binding free energies than the wildtype (Δ△G_binding_ ≤ −2.5 REU). On the other hand, for DMS based hallucinated designs (restricted at 95, 99, 100 and 100A), only 11% designs exhibit lower binding free energies while 74% exhibit binding energies identical to the wildtype (|ΔΔG_binding_| ≤ 2.5 REU; **Figure 4E**; grey bars peak at the wildtype binding energy). These calculated binding energies qualitatively align with the experimental affinity measurements from Mason *et al.*, as most of the tested designs exhibited affinities comparable to or lower than the wildtype.[46] Indeed, only one of the thirty experimentally characterized binders exhibited affinity with slight improvement over the wildtype, suggesting that a DMS restricted library may limit the potential for improving affinity over the starting/wildtype sequence.

In summary, *in silico* tests suggest that hallucination may be applied to obtain less restricted sequences of CDR H3s (and other CDRs) that can bind HER2 (and other antigens) with improved affinities.

#### Top designs exhibit favorable interface metrics

We further characterized the antibody-antigen interfaces of the screened libraries. Specifically, we calculated the shape complementarity, the number of hydrogen bonds and the total, and polar buried surface area at the antibody-antigen interface. In **Figure 4F**, we show the distribution of these metrics as a function of calculated free energy of binding. We find many designs improve interface metrics (wildtype shown as dashed lines) with a greater number of hydrogen bonds, larger buried surface area at the interface and higher shape complementarity.

## Conclusion

Antibody affinity maturation is a laborious, expensive, time-consuming, and routine task in the therapeutic antibody development pipeline. Affinity maturation is primarily centered on generating libraries of CDR regions followed by screening for expression and binding. Even a relatively short CDR H3 loop of 10 residues has a combinatorial design space of 20^10^. Such a large sequence space is difficult to sample or screen with most experimental or computational methods. As an alternative, we present a DL framework, F_v_Hallucinator, to generate sequence libraries conditioned on the structure and partial sequence of a known antibody that can be further screened for stability, affinity, and other desired properties. The F_v_Hallucinator provides a computational approach to sample the full combinatorial space available to a CDR loop only restricted by a target geometry or conformation.

With F_v_Hallucinator, we have extended the existing hallucination-based framework for protein design to the specific problem of the design of the antibody variable domain. While the previous hallucination frameworks have been aimed at designing protein scaffolds, our framework tackles the challenging task of generating highly variable subsequences for the CDR regions of antibodies that participate in antigen recognition.

On a benchmark set of 60 antibodies, for all six CDRs, the F_v_Hallucinator designs native-like CDR sequences with high sequence recovery (≥ 50%) when seeded with the wildtype sequence. The F_v_Hallucinator designs the heavy and light chain interface with mutations enriched in human repertoires.

To guide hallucination to relevant sequence spaces, we developed sequence-based losses. We demonstrated the efficacy of such restricted modes in generating sequence libraries on a large dataset of HER2 binders. In a restricted hallucination with only two positions restricted, over 50% of the designs exhibited ≥ 60% sequence identity with the binders, while a small fraction of designs exhibited 80% sequence identity with experimentally identified binders.

We further developed a pipeline that combines sequence libraries with physics-based models for screening for antigen binders. We tested our pipeline on the HER2 dataset and found that the pipeline recapitulates key residues for HER2 binding. We also show that the pipeline enables the in-silico generation of diverse screened libraries that can access significant improvements in affinity over the starting antibody.

Compared to language-based models for CDR sequence design such as IgLM[22] and the CDR manifold sampler[49], the F_v_Hallucinator pipeline enables targeted and controlled antibody subsequence design. Such a strategy could possibly lead to better and more predictable outcomes in the lab[50].

While we present comparison of hallucinated sequences with over 480 CDR loops (80 target antibodies; six CDR loops per target), PyIgClassify canonical clusters that represent distributions over all known CDR structures and an experimental CDR H3 library for the Trastuzumab antibody, further experimental verification is needed to prove that these results are useful. With the fast pace of deep-learning research in protein design, these computational findings reveal the promise of hallucination towards the fast and cheap in-silico generation of diverse, structure-conditioned antibody libraries enriched in binders. The F_v_Hallucinator framework is versatile and easily extendable to designing libraries conditioned on grafted loop conformations. Furthermore, the pipeline can be modified to screen for other engineering goals such as stability and developability.

## Methods

### Design Approach

Like Anishchenko *et al*.[12] (Hallucination) and Norn *et al*.[27] (trDesign), we aim to design sequences that fold into a desired structure using a pre-trained sequence-to-structure deep learning model. The problem of predicting sequence given structure can be stated as the problem of maximizing the probability of a sequence (*S*) given target structure (*T*). Using Bayes theorem:

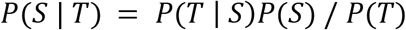

We split the sequence *S* into subsets of positions *S_D_* to be designed and *S_F_* to be fixed. To maximize *P*(*S*|*T*), we maximize the product *P*(*T*|*S*)*P*(*S*) with respect to *S_D_* with *T* and *S_F_* fixed:

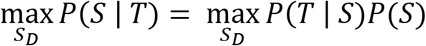

Since logarithm is a monotonically increasing function, we can apply it to equation 1b, and maximize the logarithm of *P*(*S*|*T*) to obtain Equation 1.

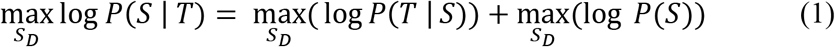

### Geometric Losses

In Equation 1, we estimate *P*(*T*|*S*) with an ensemble of pre-trained DeepAb models[28] that predict the probability of a target structure geometry (approximated by C_A_, C_B_, N, O distances and orientations).

During a hallucination run, to maximize log *P*(*T*|*S*), we minimize the categorical cross entropy loss (*L_G_*) of the pre-trained DeepAb model with respect to the design subsequence, *S_D_*. We restrict loss calculation to pairs of residues with C_A_ atoms within 10 Å of each other[25]. The cutoff value is chosen based on that used in the DeepAb study for variant prediction.[28] Thus, the objective function for minimizing geometric losses with respect to the design subsequence, *S_D_*, can be written as,

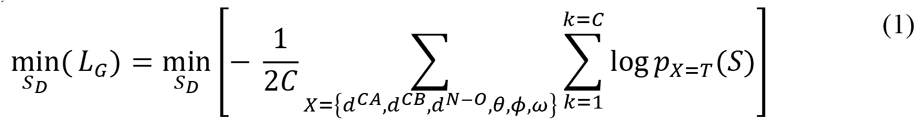

where, *p*_*x*=*T*_(*S*) is DeepAb’s predicted probability of the target geometric label *X* ∈ {*d^CA^, d^CB^, d^N-O^, θ, ϕ ω*} for sequence *S* and *C* is the number of contacts under 10 Å:

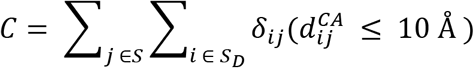

### Sequence-based losses

To maximize log(*P*(*S*)) in Equation 1, we define different priors for the sequence (*P*(*S*)) that are then reframed as sequence losses and minimized with respect to the design subsequence, *S_D_*.

#### No sequence loss

In unrestricted hallucination, we simply set *P*(*S*) = 1 in Equation 1 resulting in a sequence loss of zero.

#### Sequence loss

In sequence-restricted hallucination, we maximize *P(S)* such that categorical cross entropy loss of the designed amino acid (*z_i_*) at design position *i* is minimized with respect to the amino acid residue in the wildtype sequence at design position *i*, 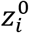.

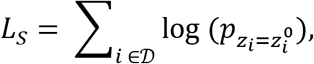

where 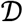 is the set of positions to be designed.

#### Motif or PSSM loss

In motif-restricted hallucination, we maximize the term *P(S)* such that the KL divergence (*D_KL_*) between the amino acid distribution of the designed sequence (*S_D_*) at the subset of motif positions *M* and the target distribution is minimized:

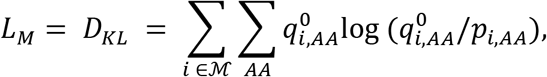

where 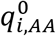 is the target/motif amino acid distribution at position *i, p_i,AA_* is hallucinated amino acid distribution at position *i*, and 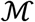 is subset of design positions at which the motif is specified.

Geometric and sequence losses may be weighted to match their magnitudes with weights *w_G_*, w_S_ and *w_M_*:

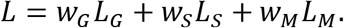

All weights are set to one by default.

### Design sequence initialization and wildtype seeding

Each element (*a_i,AA_*) of the design subsequence matrix (number of design positions x 20 amino acids) is initialized with a random number from a uniform probability distribution. When wildtype seeding is enabled, at each position *i* in the design subsequence matrix, the probability of the amino acid at the same position in the wildtype sequence, 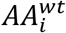, is skewed as follows:

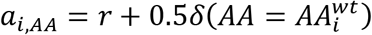

Finally, the amino acid 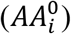 initialized at design position *i*, is the amino acid with the maximum probability at design position *i* (argmax) in the normalized (softmax) design subsequence matrix, *i.e.*,

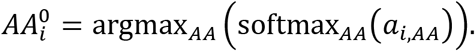

### Sequence Recovery on RAbD dataset

For sequence recovery benchmark on RAbD dataset[7], we generated 50 designs each for CDRs H1, H2, L1, L2 and L3 and 100 designs for CDR H3. Each hallucination was run with default settings (geometric loss only). Cysteine is disallowed at all positions by sampling from a reduced amino acid alphabet (see next section).

### Sampling from reduced amino acid alphabet

To sample from a reduced set of amino acids (*e.g.*, to design out cysteine or proline), we initialize from a reduced alphabet and set the gradient of the loss with respect sequence equal to zero for the unwanted amino acid residues at all design positions.

### Comparison between designed sequences and PyIgClassify clusters

We compared the distribution of designed sequences for CDR regions to the sequence profiles of PyIgClassify[8] clusters by calculating the Bhattacharya coefficient (BC) and the Bhattacharya distance (BD) at each design position. BC is a measure of the overlap between two statistical samples and the BD is a symmetric measure of the distance between two distributions.

We converted both the designed sequences and the relevant PyIgClassify cluster into PSSMs, *p* and *q* respectively. Only non-starred (usually clusters with significant number of known structures) PyIgClassify clusters were analysed. For each position *i*, we calculated the BC as 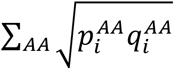 over all amino acids and BD as 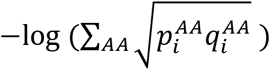. To calculate the average BC (or BD) for a CDR sequence, we averaged the BC (or BD) over all positions *i*. To avoid infinite values for BD, we replaced all zero-valued arguments in the calculation of BD with a small number (10-^6^).

For CDR H3 loops of lengths 17, 19 and higher, there is only one non-starred PyIgClassify cluster, hence, for CDR H3 targets with these lengths, no data is reported in Figure 2B and SI Figures 5-10 for the non-target clusters. For all other CDR H3 targets, there is at least one, non-starred canonical cluster available in the PyIgClassify database.

### Calculation of perplexity

The perplexity of a distribution *p*(*x*) such as each residue position of a PyIgClassify cluster sequence profile or a hallucinated sequence profile was calculated as: *PP*(*p*(*x*)) = *2*-^∑_*AAP*_(*x*)log_2_*p*(*x*)^.

### Hallucination of V_H_-V_L_ interface

For data reported in Table 1, we generated 60 hallucinated sequences per antibody without wildtype seeding. Germline gene ids were obtained with ANARCI.[51] Cysteine is disallowed at all positions by sampling from a reduced amino acid alphabet.

### Hallucination for CDR H3 of Trastuzumab

Each hallucination was run with default settings with additional sequence-based losses. Ten positions on CDR H3 (H95 – H101; Chothia numbered) were designed (to match experiments[46]) in each case. Cysteine is disallowed at all positions by sampling from a reduced amino acid alphabet. For motif-restricted hallucination, the weight for the motif-loss (*w_M_*) was set to 100. For sequence-restricted hallucination, the weight for the sequence loss (*w_S_*) was set to 25. For each hallucination run, we generated between 600 and 1,200 designs. More specifically, for data reported in Figures 3D and 3D, we generated 600 sequences for unrestricted, sequence restricted, and motif restricted (II) hallucination and 1200 sequences for the motif-restricted hallucination (I).

### Folding sequences with DeepAb

We follow Ruffolo *et al.*[28] to fold designed sequences with DeepAb.

### Calculation of free energy of binding and interface metrics with Rosetta

The free energy of binding was calculated using the InterfaceAnalyzer[48] application in PyRosetta[52]. We assumed that the designed antibody retains the wildtype binding mode *i.e.*, the epitope and paratope geometries are similar to the wildtype complex. That is, we simply threaded the designed sequences on the crystal structure of the complex and packed the side chains at the antibody-antigen interface with FastRelax. For each designed sequence, we generated 5 decoys and selected the decoy with the lowest free energy of binding (reported as dG_separated by InterfaceAnalyzer). The number of decoys (tested 2-25) did not significantly change the lowest free energy of binding.

## Availability

The full pipeline will be made available on Github upon the acceptance of the paper. Hallucinated sequences for all benchmark sets reported in the work will be deposited on Zenodo.

## Funding

This work was supported by NIH grants R01-GM078221 and R35-GM141881 and GlaxoSmithKline Vaccines. J.A.R is also supported by fellowship from AstraZeneca.

## Acknowledgements

We would like to thank Dr. Jeremias Sulam (JHU) for helpful suggestions at the early stages of the project and to Dr. Newton Wahome (GSK) for helpful discussions. We also acknowledge high performance computing resources from MARCC and ARCH.

## Author contributions

S.P.M, J.A.R, J.J.G conceptualized the work

S.P.M, J.A.R wrote the hallucination pipeline code

S.P.M wrote the original manuscript

S.P.M, J.A.R, R.F. wrote analysis scripts, tested the code, and analyzed data

S.P.M, J.A.R, R.F., J.J.G edited the manuscript

J.J.G acquired the funding

J.J.G supervised the project

## Competing interests

J.J.G is an unpaid board member of the Rosetta Commons. Under institutional participation agreements between the University of Washington, acting on behalf of the Rosetta Commons, Johns Hopkins University may be entitled to a portion of revenue received on licensing Rosetta software including methods discussed/developed in this study. J.J.G has a financial interest in Cyrus Biotechnology. Cyrus Biotechnology distributes the Rosetta software, which may include methods developed in this study. These arrangements have been reviewed and approved by the Johns Hopkins University in accordance with its conflict-of-interest policies.

## Notes

### Summary of Updates

Slight changes in wording in the abstract and introduction to emphasize targeted library generation. Also, added Peter Kim's citation (Hie et al.).

