## SupplementaryFigures for "Hallucinating structure-conditioned antibody libraries for target-specific binders"

### Supplementary Figures

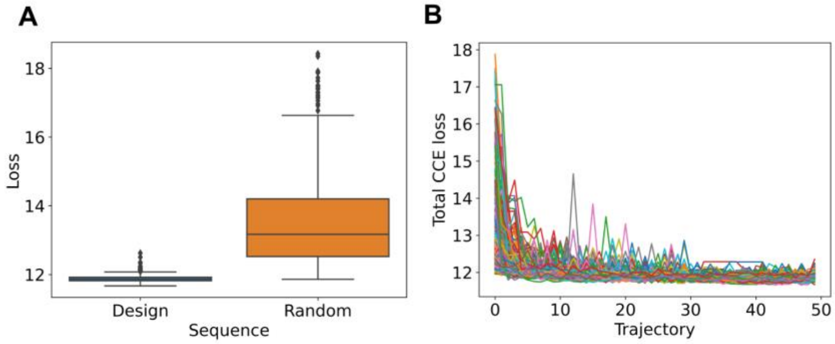


**SI Figure 1. Hallucination minimizes geometric losses of designed sequences. A. Comparison of distribution of CCE loss of the designed sequences versus random sequences they were initialized with. B. Minimization of CCE losses over ~200 hallucination trajectories for an example CDR H3 loop.**

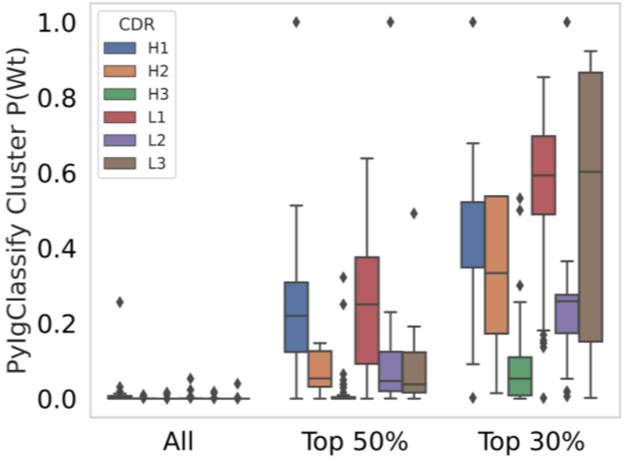


**SI Figure 2. Likelihood of recovering wildtype sequence based on PyIgClassify sequence profile for cluster corresponding to wildtype sequence for "ALL" positions (or 100% sequence recovery) on the CDR, for "Top 50%" most conserved positions and for "Top 30%" most conserved positions.**

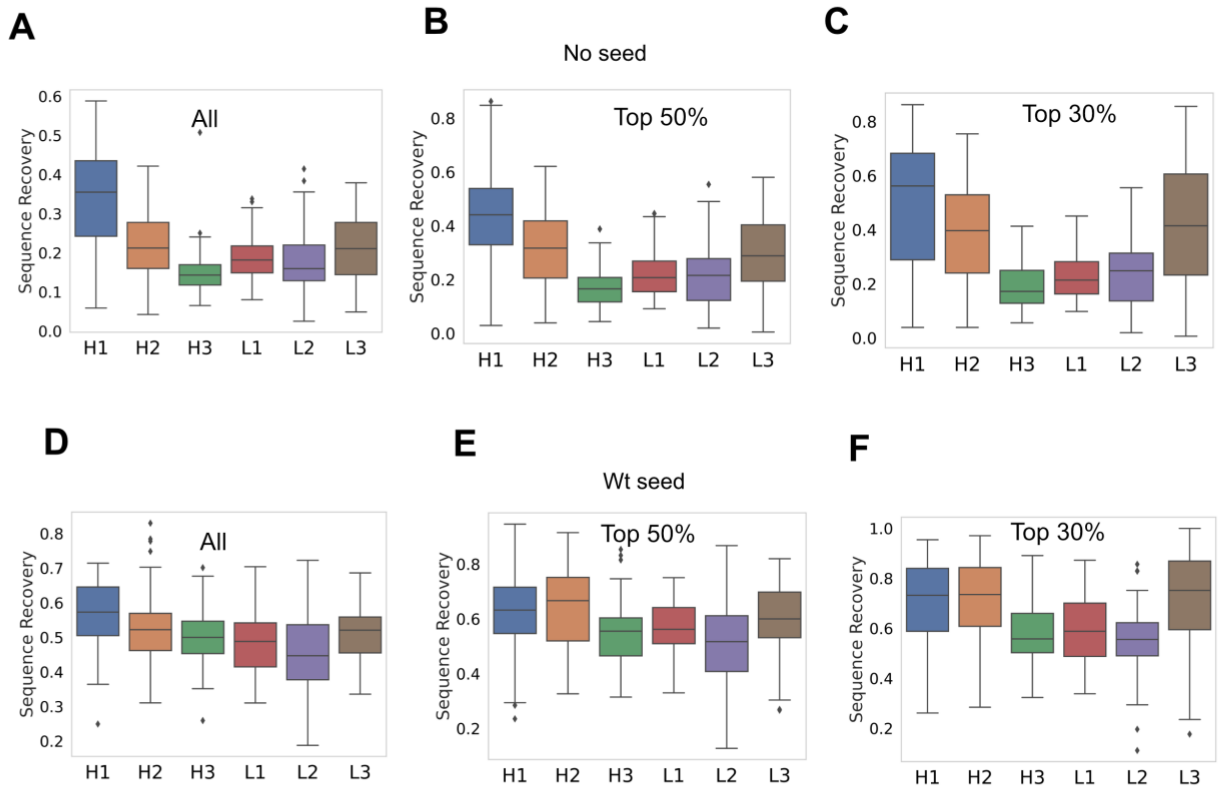


**SI Figure 3. Sequence recovery for "All" positions, "Top 50%" most conserved positions and "Top 30%" most conserved positions (conservation estimated from PyIgClassify cluster profiles) for hallucination without seeding (A, B, C) and with seeding (D, E, F).**

| PDB IDs |
| --- |

| |
|---|
| 1bey, 1cz8, 1dlf, 1fns, 1gig, 1jfq, 1jpt, 1mfa, 1mim, 1mlb, 1mqk, 1nlb, 1oaq, 1seq, 1sy6, 1yy8, 2d7t, 2e27, 2fb4, 2fbj |

**SI Table 1. PDB ids of 20 antibodies selected from DeepAb Test set[1] for sequence recovery.**

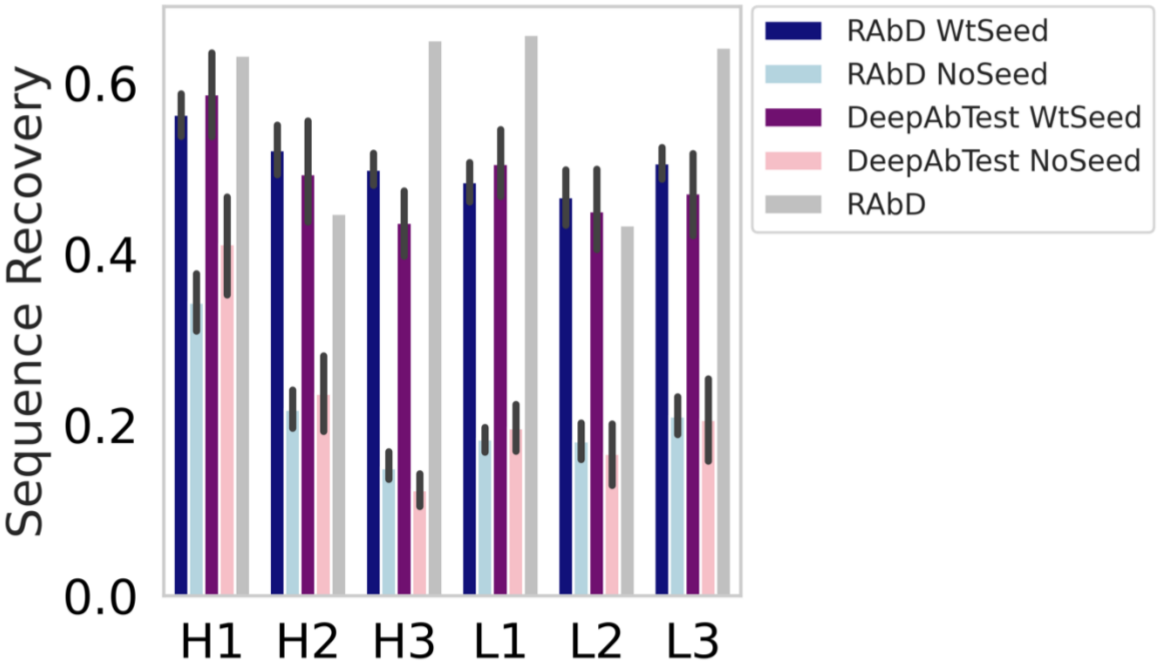


**SI Figure 4. Sequence recovery with (DeepAbTest WtSeed) and without wildtype seeding (DeepAbTest NoSeed) on a benchmark set of 20 antibodies selected from the DeepAb test set. This benchmark set was "blind" to the pre-trained DeepAb model used in this work. The sequence recovery numbers are similar to those obtained for the RAbD dataset with and without wildtype seeding. The 54/60 antibodies in the RAbD dataset were also in DeepAb's training set. However, no difference is sequence recovery is observed between the DeepAb Test set and the RAbD dataset. We have also included the sequence recovery of the RAbD method on the RAbD dataset (RAbD) for comparison. See SI Table 1 for the pdb ids of 20 antibodies chosen from the DeepAb test set.**

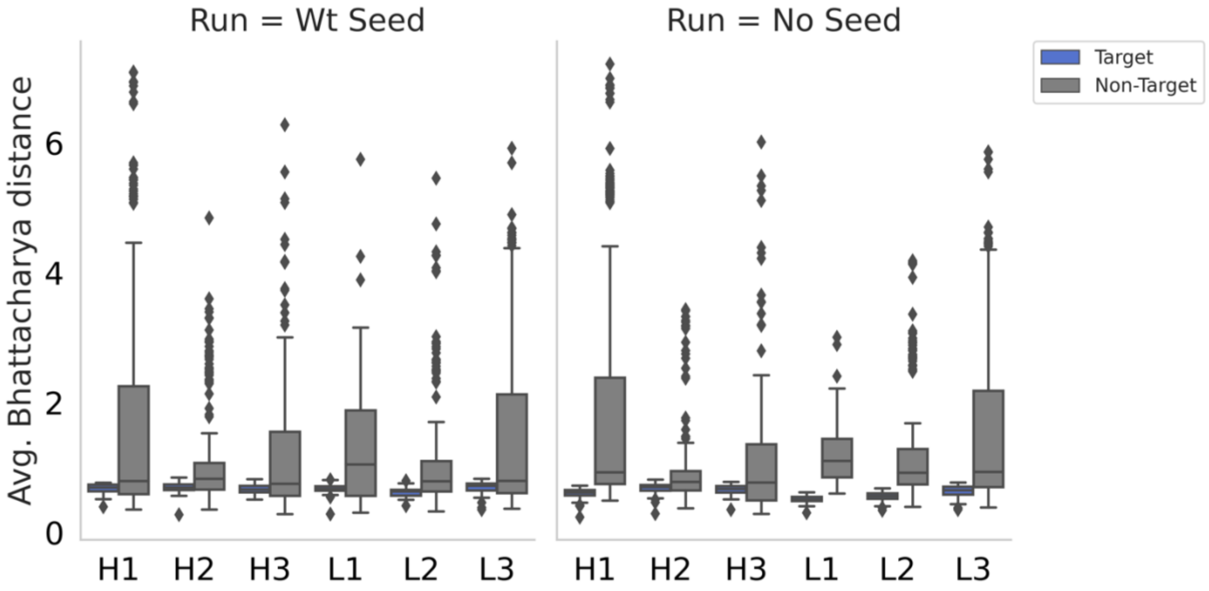


**SI Figure 5. Average Bhattacharya distance (BD) between the hallucinated sequence and the Target and non-Target PyIgClassify clusters. BD is the negative logarithm of BC and a measure of the distance between two distributions. BD is first averaged over all hallucinated sequences for each target and then averaged over all targets for each CDR. Sequence profiles of hallucinated designs are closer to the target (blue) PyIgCluster than non-target (gray) clusters.**

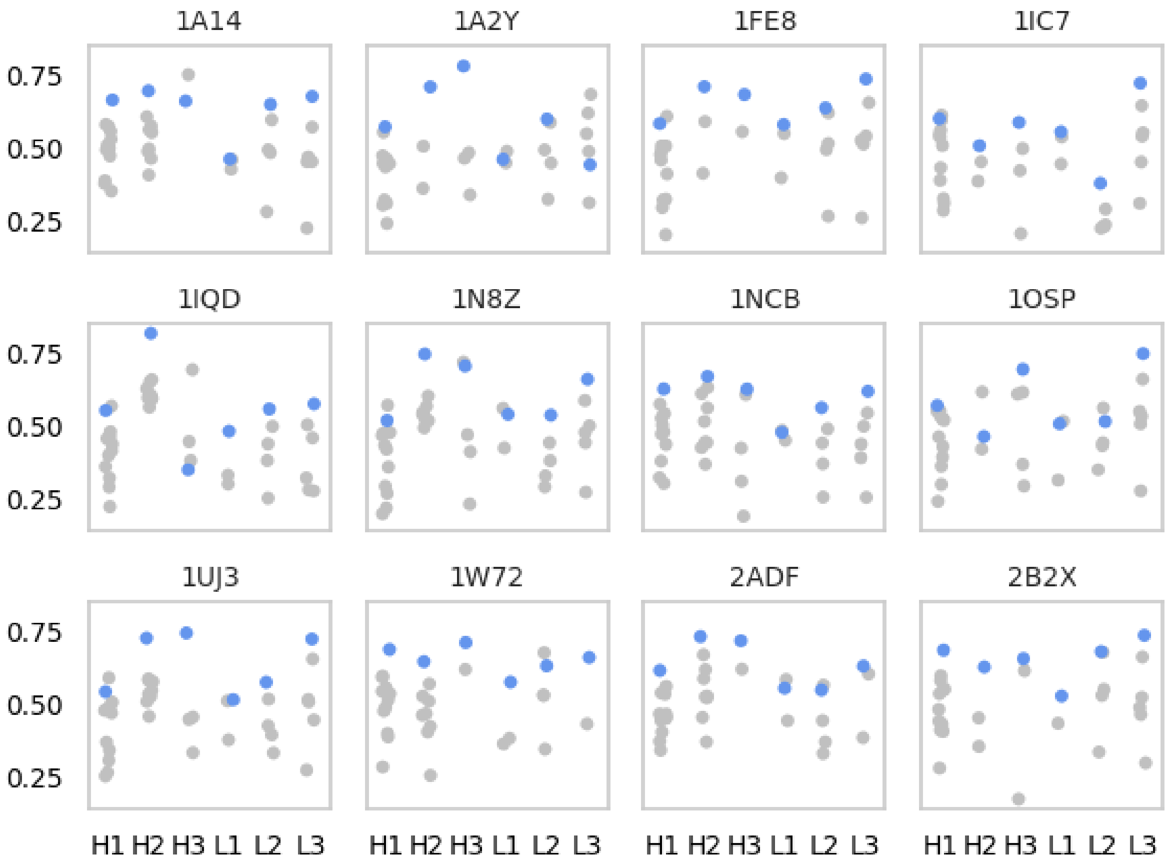


**SI Figure 6. Bhattacharya coefficient averaged over all designed positions between the sequence profile from 50 hallucinated (no wildtype seeding) CDRs and the *target cluster (blue)* and non-target cluster (gray) for all 6 CDRs for targets 1-12 from the RAbD benchmark set.**

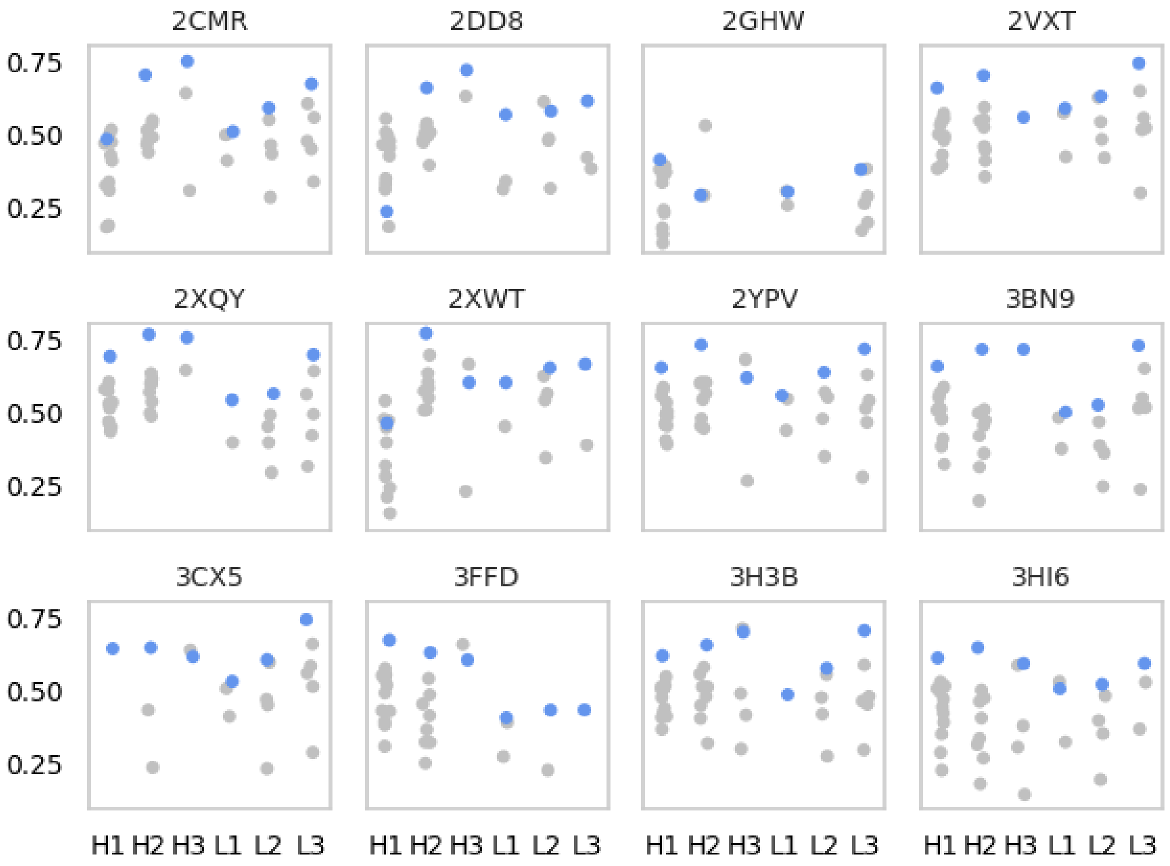


**SI Figure 7. Bhattacharya coefficient averaged over all designed positions between the sequence profile from 50 hallucinated (no wildtype seeding) CDRs and the *target cluster (blue)* and non-target cluster (gray) for all 6 CDRs for targets 12-24 from the RAbD benchmark set. For 2GHW H3, the PyIgClassify target clusters are "starred" clusters. These clusters are considered unreliable and not considered in our analysis.**

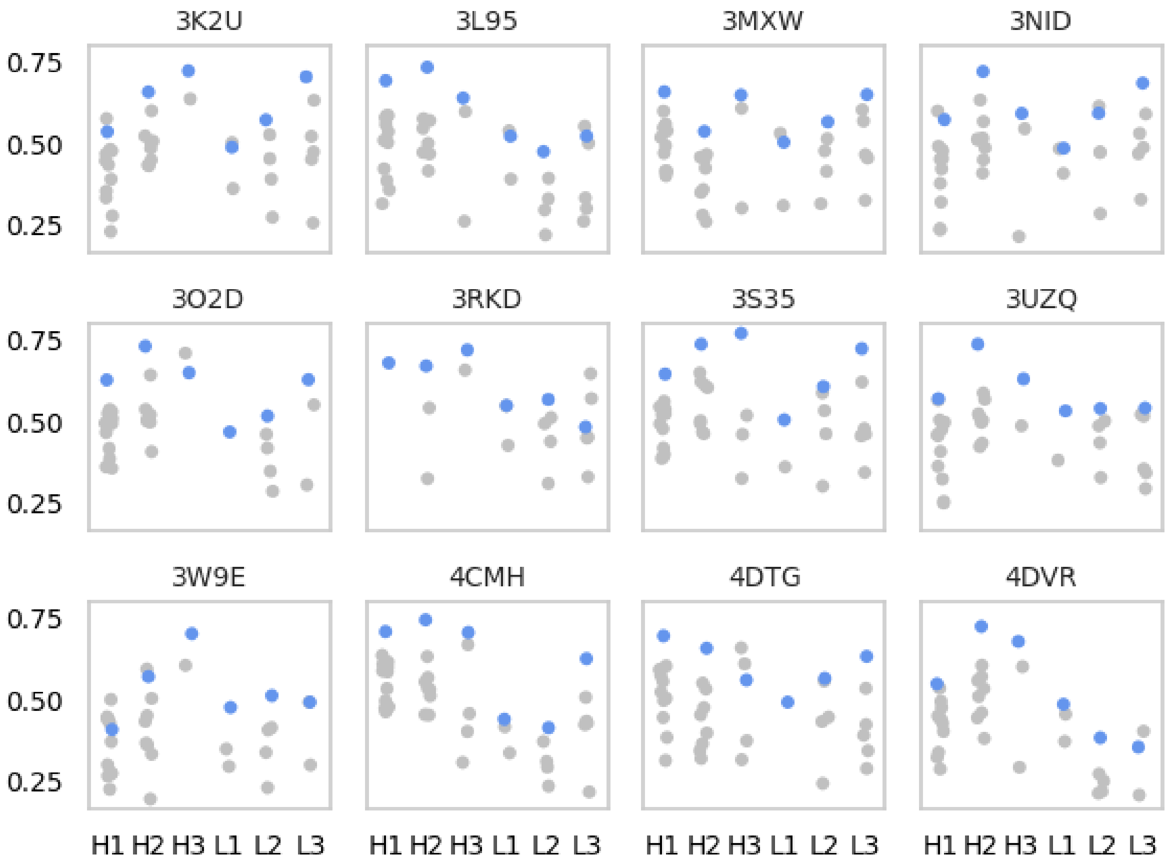


**SI Figure 8. Bhattacharya coefficient averaged over all designed positions between the sequence profile from 50 hallucinated (no wildtype seeding) CDRs and the *target cluster (blue)* and non-target cluster (gray) for all 6 CDRs for targets 24-36 from the RAbD benchmark set.**

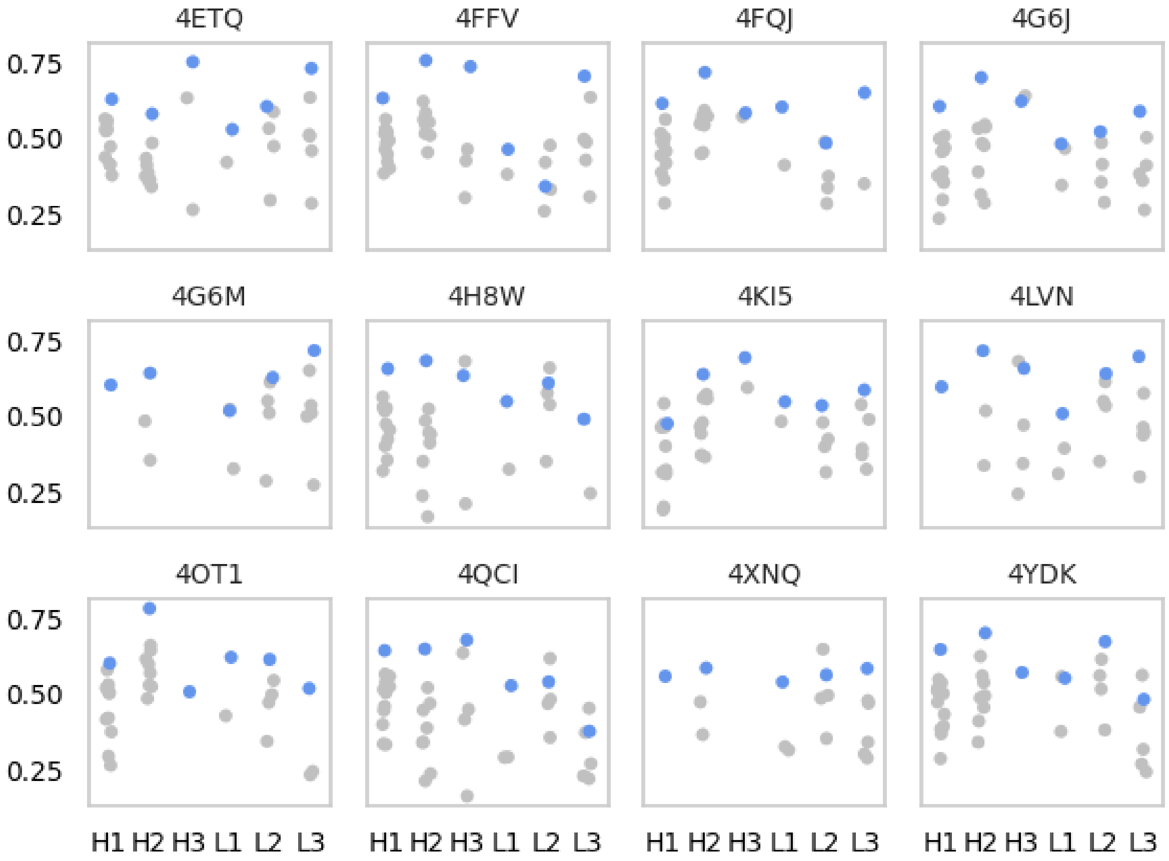


**SI Figure 9. Bhattacharya coefficient averaged over all designed positions between the sequence profile from 50 hallucinated (no wildtype seeding) CDRs and the *target cluster (blue)* and non-target cluster (gray) for all 6 CDRs for targets 26-48 from the RAbD benchmark set. For 6G6M H3 and 4XNQ H3, the PyIgClassify target clusters are "starred" clusters. These clusters are considered unreliable and not considered in our analysis.**

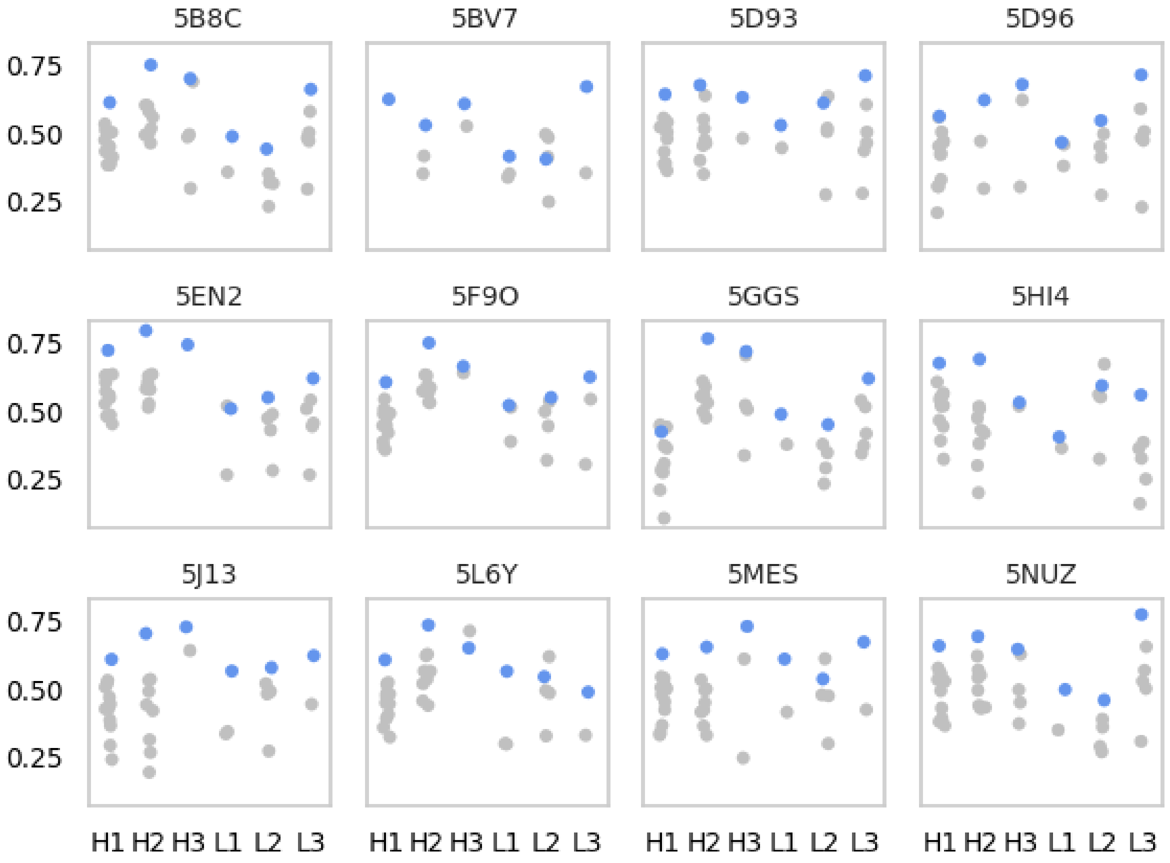


**SI Figure 10. Bhattacharya coefficient averaged over all designed positions between the sequence profile from 50 hallucinated (no wildtype seeding) CDRs and the *target cluster (blue)* and non-target cluster (gray) for all 6 CDRs for targets 48-60 from the RAbD benchmark set.**

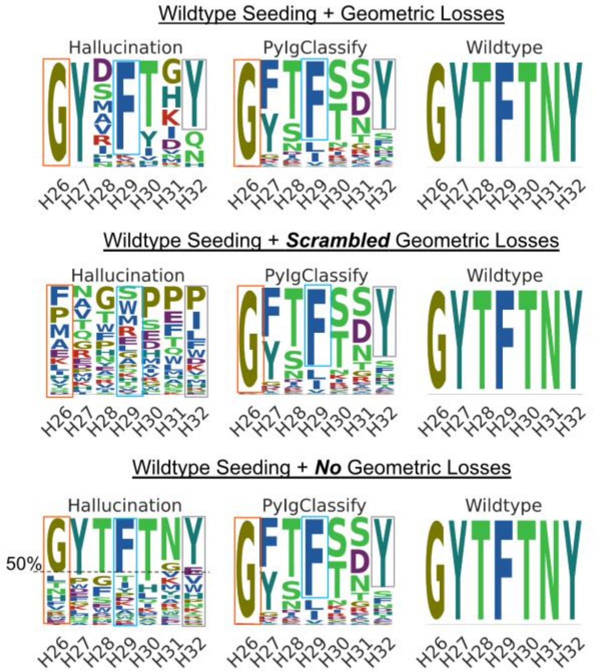


**SI Figure 11. Effect of wildtype seeding and geometric losses on sampled profiles. Geometric losses guide hallucination to sample profiles resembling those of target structural clusters. When geometric losses are scrambled, wildtype seeding does not result in either retention of the wildtype sequence nor sampling of PyIgClassify target cluster-like profiles. Thus, wildtype seeding with incorrect geometric losses generates random sampling. When geometric losses are completely removed, the wildtype sequence is sampled about 50% of the times for all positions same as the degree to which the wildtype sequence is spiked in the initialization of the designed sequence. However, the sequence profiles do not resemble those observed for PyIgClassify clusters. For example, *with geometric losses (top panel)*, H26 almost always samples a glycine same as PyIgClassify. This is not true when geometric losses are absent.**

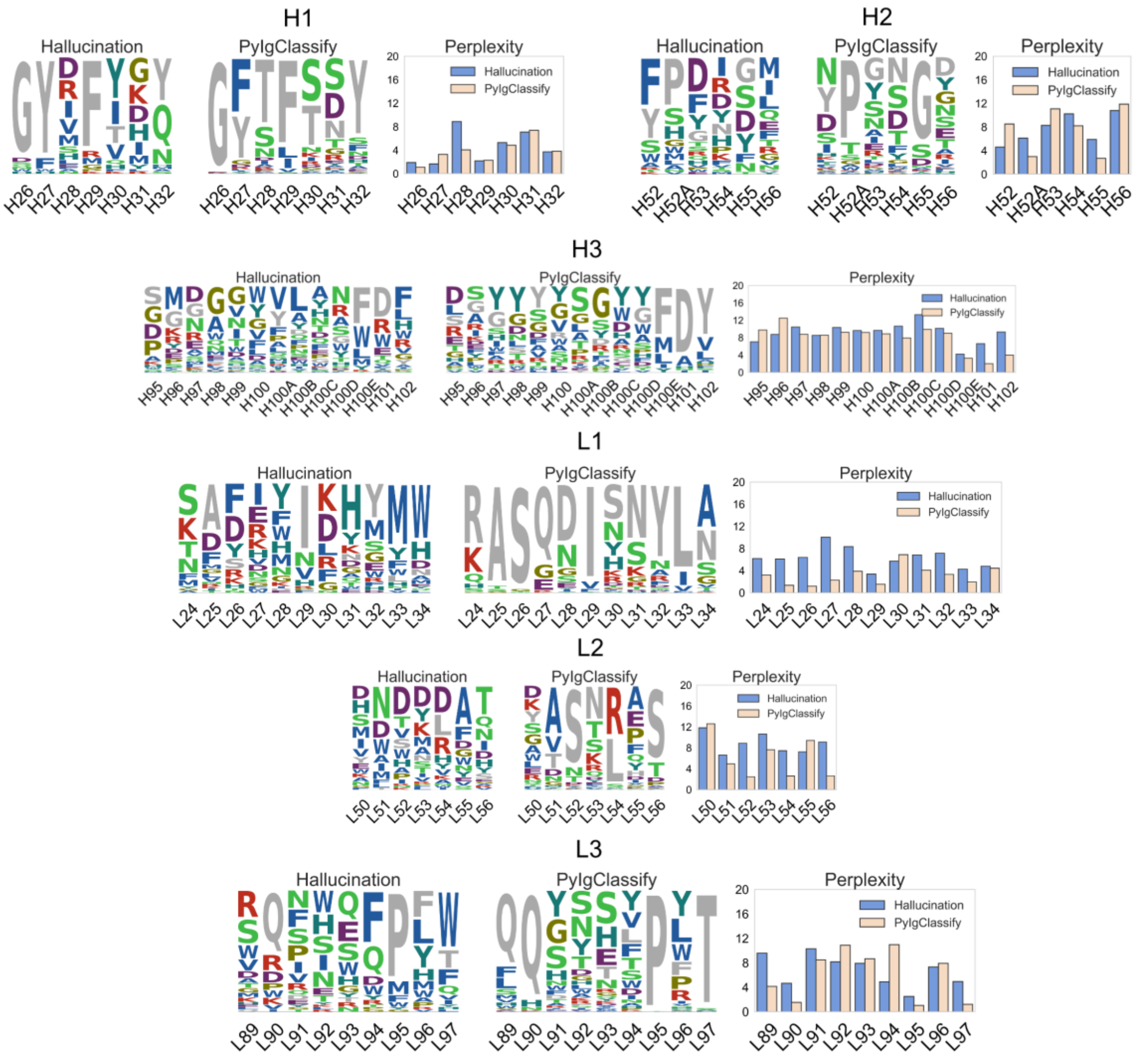


**SI Figure 12. Comparison of perplexity of per-residue probability distribution derived from hallucinated designs (without wildtype seeding) for CDRs of antibody (PDB: 1A14, RAbD dataset) and the target PyIgClassify cluster. Wildtype sequence is shown in grey. Each sequence logo was constructed from 50 designs for CDRs H1, H2, L1, L2 and L3 and 100 designs for CDR H3.**

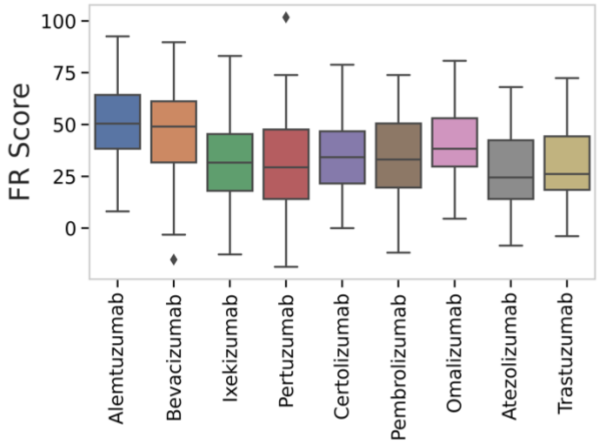


**SI Figure 13. FR scores per design for hallucinated designs for the $V_H$-$V_L$ interface for the humanized antibodies dataset.**

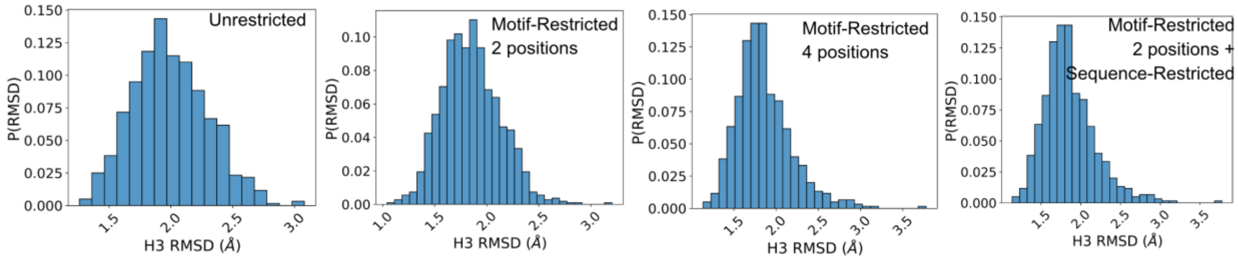


**SI Figure 14. Distribution of H3 RMSD of the forward folded designs from different hallucination runs for the Trastuzumab antibody. For all modes shown here except unrestricted hallucination, >70% of hallucinated sequences have an H3 RMSD of under 2 Å from the target CDR H3. For unrestricted hallucination, only 326 of 600 designs (54%) have an H3 RMSD of under 2 Å from the target CDR H3.**

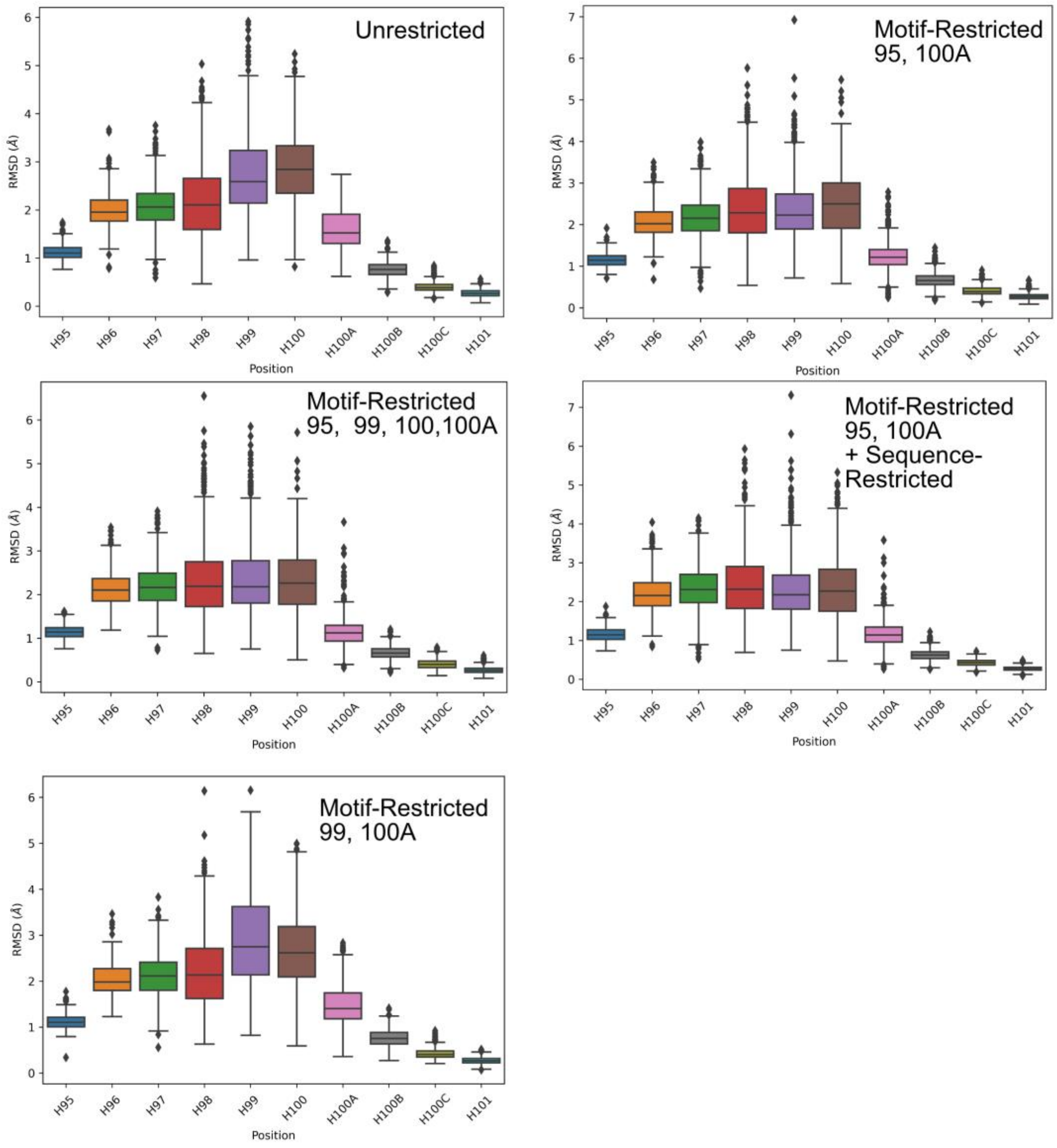


**SI Figure 15. Distribution of per-residue backbone heavy atom RMSDs of forward folded designs from different hallucination runs for the Trastuzumab antibody**

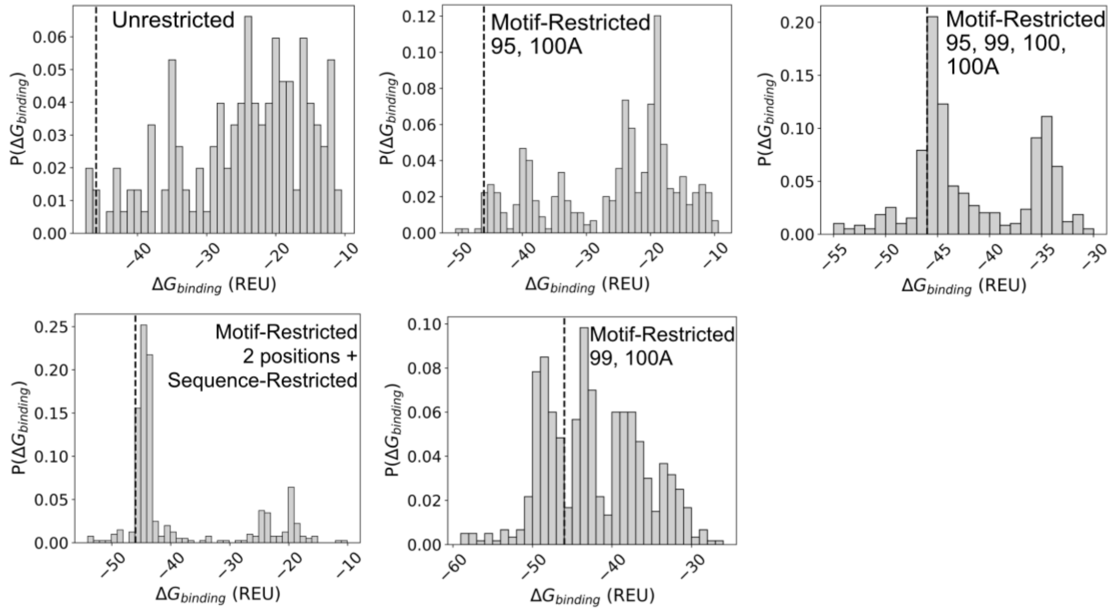


**SI Figure 16. Distribution of $\Delta G_{binding}$ between HER2 and designs from different hallucination runs for the Trastuzumab antibody. Wildtype $\Delta G$ is indicated by the dashed line.**

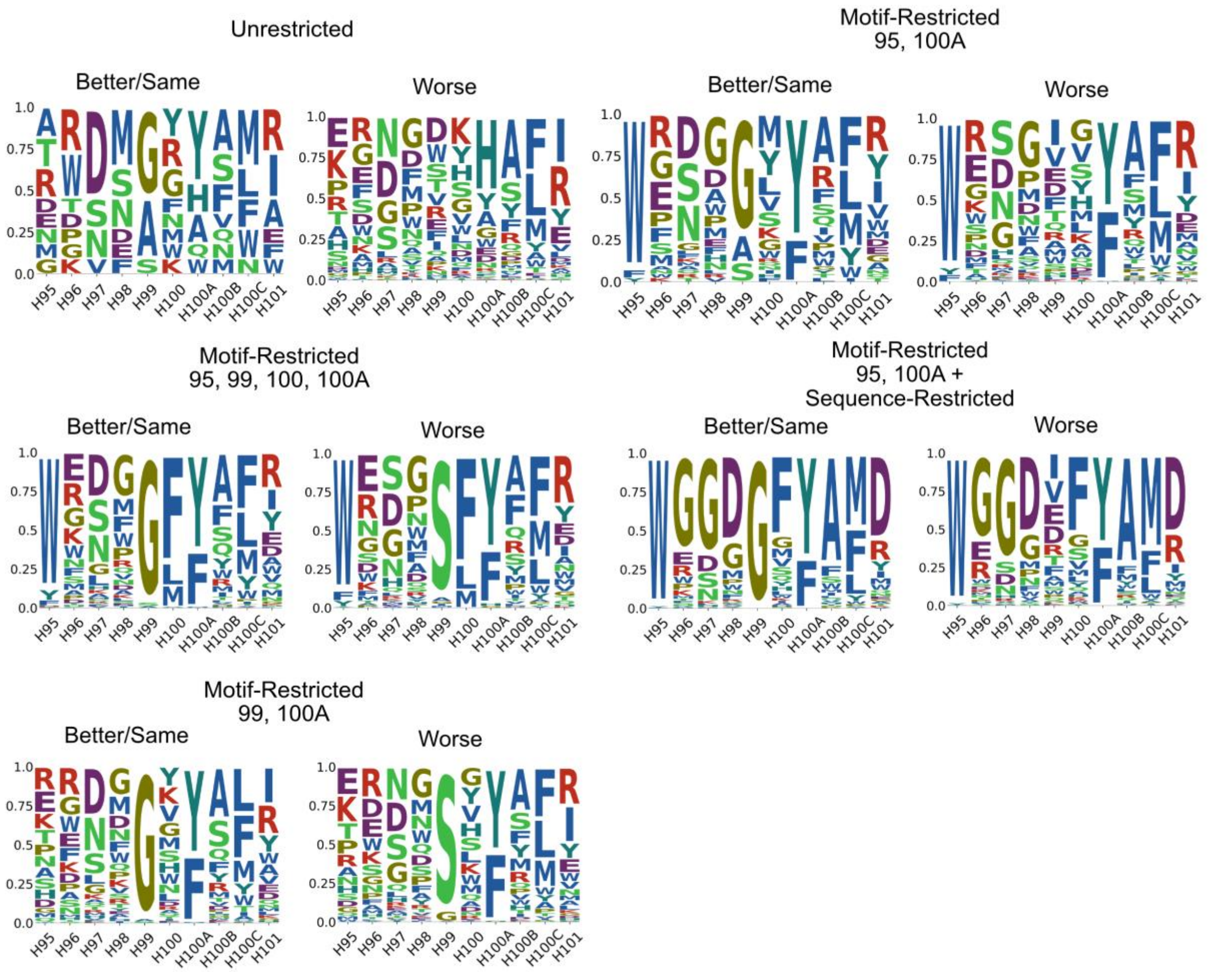


**SI Figure 17. Sequence logos for designs with binding free energy either "Better/Same" as the wildtype or "Worse" than the wildtype after virtual screening for hallucinated libraries in different hallucination modes for the Trastuzumab antibody CDR H3.**

(1) Ruffolo, J. A.; Sulam, J.; Gray, J. J. Antibody Structure Prediction Using Interpretable Deep Learning. *Patterns* **2022**, *3* (2), 100406. https://doi.org/10.1016/j.patter.2021.100406.
